# Immunomagnetic Sequential Ultrafiltration (iSUF) Platform for Enrichment and Purification of Extracellular Vesicles from Biofluids

**DOI:** 10.1101/2020.05.13.089573

**Authors:** Jingjing Zhang, Luong TH Nguyen, Richard Hickey, Nicole Walters, Xinyu Wang, Kwang Joo Kwak, L. James Lee, Andre F. Palmer, Eduardo Reátegui

## Abstract

Extracellular vesicles (EVs) derived from tumor cells have the potential to provide a much-needed source of non-invasive molecular biomarkers for liquid biopsies. However, current methods for EV isolation have limited specificity towards tumor-derived EVs that limit their clinical use. Here, we present an approach called immunomagnetic sequential ultrafiltration (iSUF) that consists of sequential stages of purification and enrichment of EVs (nonspecifically and specifically) in approximately 2 h. In iSUF, EVs present in different volumes of biofluids (0.5 mL to 100 mL) can be significantly enriched (up to 1000 times), with up to 99 % removal of contaminating proteins (e.g., albumin). The EV recovery rate by iSUF for cell culture media (CCM), serum, and urine corresponded to 98.0% ± 3.6%, 96.0% ± 2.0% and 94.0% ± 1.9%, respectively (p > 0.05). The final step of iSUF enables the separation of tumor-specific EVs by incorporating immunomagnetic beads specific to a target subpopulation of EVs. Serum from a small cohort of clinical samples from metastatic breast cancer (BC) patients and healthy donors were processed by the iSUF platform and the isolated EVs from patients showed significantly higher expression levels of BC biomarkers (i.e., HER2, CD24, and miR21).

## Introduction

Extracellular vesicles (EVs) are increasingly recognized as relevant diagnostic and therapeutic entities present in different biofluids^1^. EVs are lipid particles with sizes that vary from 30 nm to a few microns^2^. EVs are endogenously shed from the surface of cells through distinct mechanisms, leading to different types of vesicles^3^. Multivesicular bodies that contain smaller vesicles can fuse with the plasma membrane to release their internal vesicles (i.e., exosomes)^4^. Larger lipid vesicles can directly bud from the plasma membrane as microvesicles^5^. EVs carry various biological cargo, including proteins, RNA, and DNA fragments, giving EVs unique roles in regulating cell-cell communication^6^. Moreover, it has been shown that tumor EVs (tEVs) can tune cellular microenvironments at distant sites to promote angiogenesis, invasiveness, immunosuppression, and metastasis^7–10^.

Different proof of concept studies have used tEVs to develop liquid biopsy assays to diagnose and monitor cancer at different stages^11,12^. EVs are more abundant than other circulating biomarkers (e.g., circulating tumor cells), and they are structurally more robust^13^. However, tEVs present in biofluids are surrounded by massive amounts of normal EVs (nEVs; secreted by healthy cells), and other biomolecules (e.g., albumin, lipoproteins, globulins)^14^, thus novel purification methods are required to isolate tEVs^15^. A recent survey on the methods used for isolation and characterization of EVs from research laboratories around the world reveals that more than 80% of researchers use ultracentrifugation (UC) for the isolation of EVs and western blotting for protein characterization^16^. Although UC and density gradient methods can be used to process different biofluids, they are labor-intensive, produce protein aggregate contaminants, and are nonspecific towards EV type (derived from a tumor or normal cells)^17–19^. Other EV isolation methods, including polymeric or salt precipitation kits^20^, size exclusion chromatography (SEC) columns (e.g., qEVs)^21^, and nano/microdevices have limitations^22^. Precipitation kits have low EV recovery rates, lack specificity, and have low purity^23^. qEVs can separate EVs into different size fractions with high purity and low protein contamination, but have low EV recovery rates and are nonspecific for EV subpopulation^24^. Recently, immunoaffinity methods that were developed for cell separation have been adapted for specific EV isolation^25^. Microfluidic and plasmonic devices have been functionalized with antibodies to target different EV populations^26,27^. However, the majority of these approaches target tetraspanins and annexins, which are ubiquitous proteins present in all EVs^28^. Other attempts used epithelial cell adhesion molecule (EpCAM); however, this antigen is also expressed on normal epithelial EVs^29^. Recently, we demonstrated the use of nanostructured polymeric brushes conjugated with epidermal growth factor receptor (EGFR) and integrated into a microfluidic channel to enhance specificity towards tumor-derived EVs isolated from glioblastoma (GBM) patients^30^. Although this approach can achieve a remarkable 94% specificity towards tEVs, the limited amount of biofluid processed (1 to 1.5 mL of serum or plasma) and the retention of albumin, significantly limits its use for proteomics^31^.

Compromises have to be made when using a particular technology/methodology for the isolation of EVs^32^. Currently, there is a trade-off between sample volume and specificity in EV isolation technologies that limits quantitative molecular analysis of EV contents, ultimately impacting the utility of EVs in cancer diagnostics^33^. Here, we present a novel approach termed immunomagnetic sequential ultrafiltration (iSUF) that overcomes current limitations for EV enrichment and purification. iSUF combines three stages of ultrafiltration and immunoaffinity separation: a tangential flow filtration (TFF) step, a standard centrifugal enrichment step, and a magnetic-bead antibody-based EV capture step. Using iSUF, we demonstrate that small or large volumes of biofluid can be processed (~ 500 μL or > 100 mL) while concomitantly removing up to 99 % of contaminating proteins (e.g., albumin, lipoproteins, globulins). We have demonstrated the use of iSUF for the enrichment of EVs present in three different types of biofluids: cell culture media (CCM), serum, and urine for which the sample processing time was under approximately 2 h. Another feature of iSUF is that it can enrich EVs up to 1000 times with an EV recovery rate higher than 94 %, which overcomes the limitations of other commercially available methods. To further validate the clinical utility of iSUF, we have processed serum samples from 10 metastatic breast cancer (BC) patients and demonstrated the presence of HER2, CD24 and miR21 biomarkers at significantly higher levels compared to healthy controls (p < 0.05).

## Materials and Methods

### Ethics

Healthy donors (HDs) were enrolled via an approved Institutional Review Board at The Ohio State University (IRB# 2018H0268 and 2019C0189). All HDs participants provided written informed consent. We also confirm that all experiments were carried out following relevant guidelines and regulations.

### Materials

Hollow fiber cartridges (molecular weight cut off, MWCO: 500 kDa, material: polysulfone) were supplied by Repligen (Rancho Dominguez, CA). Amicon^®^ ultra-15 centrifugal filter units (MWCO: 3, 10, and 30 kDa) were purchased from MilliporeSigma (Burlington, MA). Streptavidin-coated magnetic particles (3.0-3.9 μm) were obtained from Spherotech (Lake Forest, IL). For capturing EVs, Cetuximab was purchased from ImClone LLC (Branchburg, NJ), EpCAM, CD63, HER2 were obtained from R&D Systems (Minneapolis, MN). Antibodies were biotinylated using an EZ-Link™ micro Sulfo-NHS-biotinylation kit (ThermoFisher Scientific, Waltham, MA). For surface marker EV detection, CD24 was obtained from Novus Biological LLC (Centennial, CO), and HER2 was obtained from Cell Signaling Technology (Danvers, MA).

### Cell culture and supernatant collection

U-251 glioblastoma (GBM), MCF-7 breast, and A375 melanoma cancer cell lines were supplied by American Type Culture Collection (ATCC, Manassas, VA). Cell lines were cultured in their recommended culture medium^34^ containing 10% FBS and 1% penicillin-streptomycin at 37°C in a 5% CO_2_ incubator. For isolation of EVs from CCM, U251, MCF7, and A375 were grown in T75 flasks to 90% cell confluence, followed by washing the cells twice with PBS. Culture medium with 10 % EV-depleted FBS was added to cells for 24 h. CCM was centrifuged at 1, 000 x *g* for 5 min at room temperature (RT) to discard cell debris before further processing. EV-depleted FBS was prepared by using the permeate of FBS filtered by tangential flow filtration (TFF) (MWCO: 300 kDa).

### Healthy donor serum collection

10 mL of whole blood from healthy donors was collected into BD SST serum tubes (Thermo Fisher Scientific, Waltham, MA). Tubes were rocked 10 times and then gently placed upright to coagulate for 60 min. Then, the tubes were centrifuged at RT at 1,100 × *g* for 10 min. The serum was subsequently aspirated carefully and stored in 1 mL aliquots at −80 °C. Additionally, we purchased six vials of healthy donor serum samples from ZenBio (Research Triangle Park, NC) collected according to FDA guidelines.

### Healthy donor urine collection

Urine was also collected from the same healthy donors above, by either a first-morning or second-morning standard collection protocol^35^. The urine volume collected was 10 to 100 mL. Urine was collected in sterilized 50-mL centrifuge tubes containing 4.2 mL protease inhibitor - a mixture of 1.67 mL 100 mM sodium azide (NaN_3_), 2.5 mL phenylmethylsulfonyl fluoride (PMSF), and 50 μl Leupeptin (Millipore Sigma)^36^. After collection, urine samples were frozen at −80 °C until processing time.

### Cancer patient samples

1 mL of serum was collected from 10 metastatic BC patients. Samples were stored at −80 °C until use. All patient samples were collected from the biospecimens biobank through the Total Cancer Care (TCC) Program at the James Comprehensive Cancer Center at The Ohio State University.

### Processing biofluids using the iSUF platform

The schematic workflow of the iSUF platform is shown in **Fig. 1**. In stage 1, TFF was used to enrich and diafiltrate EVs from the biofluid. Thus, biofluid from the sample feed reservoir was removed as filtrate/permeate from the TFF filter. Diafiltration is a fractionation process that removes smaller molecules (filtrate/permeate) through the filter and leaves larger molecules in the reservoir by adding a diafiltration solution into the reservoir at the same rate as the filtrate is generated. Briefly, a TFF pump circulates the biofluid through a hollow fiber filter cartridge at a controlled flow rate. Sample fractionation depends on the hollow fiber membrane pore size (MWCO), which should be large enough to permeate proteins and free nucleic acids while small enough to retain EVs. During the enrichment step, freely permeable molecules are partially removed. To remove the remaining contaminants, a diafiltration step with PBS is necessary. The diafiltration processing time is proportional to the biofluid volume in the system^37^, so diafiltration started with a total biofluid volume of 7 mL, which was the sum of the dead volume of the self-build TFF system (2 mL in the product container) and 5 mL remaining in the tubing that can be subject to further processing later. The liquid remaining in the tubing was necessary to protect the EVs from drying out and to enable constant volume diafiltration. Hence, CCM and urine were pre-enriched to a total volume of 7 mL. For the processing of serum, 0.5 mL of sample was diluted to a total volume of 7 mL in PBS and then processed with diafiltration. The input flow rate was kept at 35 mL/min using a peristaltic pump (Cole-Parmer, Vernon Hills, IL). The sample volume after stage 1 was approximately 2 mL (dead volume of the TFF system). At stage 2, ultra-centrifugal units with 10 kDa MWCO were used to further enrich the samples to 100 μL at 3,000 × *g* for 20 min. For specific isolation of subpopulations of EVs, stage 3 of iSUF, Streptavidin-coated magnetic beads were functionalized with biotinylated capture antibodies (e.g., EpCAM, HER2, EGFR) overnight at 4 °C to target tEVs in spiked samples as well as patient samples. 100 μL of the processed sample (after stage 2) was incubated with the antibody-coated beads for 1 h at RT. The efficiency of the iSUF platform for isolating tEVs was evaluated using flow cytometry and fluorescence microscopy. EVs were also characterized for their size, concentration, morphology, and molecular content (e.g., protein, RNA).

**Figure 1.**
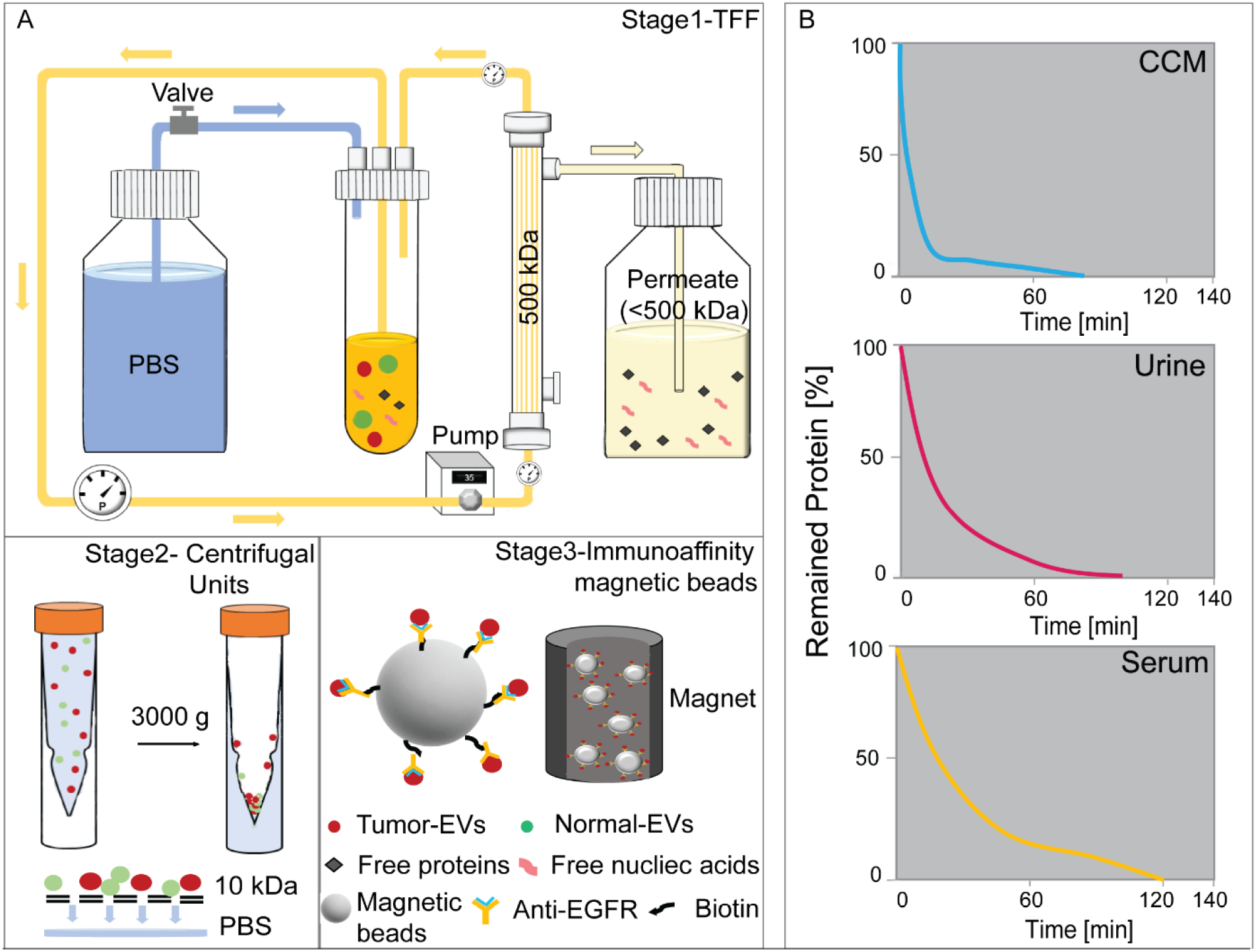
The Immunomagnetic sequential ultrafiltration (iSUF) platform. A) Schematic representation of iSUF stages 1, 2, 3, including tangential flow filtration (TFF) purification, centrifugal unit enrichment, and tEV immunoaffinity isolation, respectively. In stage 1, biofluids were processed using a 500 kDa TFF filter, EVs were retained in the retentate and enriched in 7 mL while free proteins and nucleic acids permeated through the TFF filter. Then the PBS valve was opened to start TFF diafiltration until removing up to 99 % of free proteins. Finally, EVs were recovered in 2 mL of PBS after flushing the TFF system with air. In stage 2, EVs were centrifuged using a 10 kDa centrifugal unit at 3,000 x *g* and enriched in 100 μL. In stage 3, tEVs were captured using antibodies immobilized to streptavidin-coated magnetic beads (i.e., EGFR) and subsequently pulled out with a magnet. B) iSUF stage 1 purification performance for different biofluids. Cell supernatant (CCM), urine, and serum took 80, 100, and 120 min to obtain up to 99 % efficiency of free protein and nucleic acid removal.

### Processing biofluids using ultracentrifugation (UC)

CCM, serum, and urine samples were filtered using a syringe filter (pore size: 1.0 μm) and transferred to ultracentrifuge tubes (Beckman Coulter, Brea, CA) gently using a syringe and blunt needle (Becton, Dickinson and Company, Franklin Lakes, NJ). Ultracentrifuge tubes were sealed with a cordless tube topper (Beckman Coulter) after balancing, then were placed in a Type 55.2 Ti rotor (Beckman Coulter) and centrifuged in the Optima L-80 XP ultracentrifuge (Beckman Coulter) for 90 min at 4 °C at 100,000 × *g*. The supernatants were discarded carefully after UC, and pellets were re-suspended in 100 μL of PBS.

### Processing biofluids using commercially available EV isolation kits

Using the Total Exosome Isolation Reagent (TEIR, Invitrogen, Carlsbad, CA), EVs were isolated from 0.5 mL serum according to the manufacturer’s instructions. Briefly, 0.5 mL of serum was mixed with a proprietary reagent provided in a kit and incubated for 30 min at 4 °C. After mixing, the sample was centrifuged at 10,000 × *g* for 10 min at RT. EVs pellets were resuspended in 100 μL PBS. For size-exclusion chromatography, 0.5 mL serum was loaded into a qEV column (Izon Science, Medford, MA), and flushed with PBS, fractions 7-12 were collected and pooled in 3 mL according to the manufacturer’s instructions.

### EV quantification

The size and concentration of EVs were measured using a tunable resistive pulse sensing (TRPS) method (qNano, Izon Science). Different nanopore stretchable membranes (NP150, NP300, NP600, NP800, and NP1000) were selected to cover the wide size range of EVs^38^. Samples were first filtered through 1-μm filters before processed in the qNano instrument. EVs with a size range of 70 nm to 1000 nm were characterized.

### Immunofluorescence staining

After EVs were captured on the functionalized magnetic beads using a cocktail of a recombinant chimeric EGFR monoclonal antibody (Cetuximab, Erbitux^®^, ImClone LLC, Branchburg, NJ), a goat EpCAM/TROP-1 polyclonal antibody (#AF960, R&D Systems, Minneapolis, MN) and a goat ErbB2/Her2 polyclonal antibody (#AF1129, R&D Systems), they were blocked with 3% (w/v) BSA and 0.05% (v/v) Tween^®^ 20 in PBS for 1 h at RT, and then stained with either a rabbit HER2/ErbB2 monoclonal antibody – PE conjugate (#98710S, Cell Signaling Technology, Danvers, MA) or a mouse CD24 monoclonal antibody - Alexa Fluor™ 594 conjugate (#NB10077903AF594, Novus Biologicals™) for 1 h at RT.

### Molecular beacon design and quantification

Molecular beacon (MB) (listed 5’ - 3’) targeting miR-21 used in this study was T+CA A+CA /iCy3/ +TCA +GT+C T+GA TAA GCT AAC TTA TCA GAC TGA /3BHQ_2/. Locked nucleic acid (LNA) nucleotides (positive sign (+) bases) were incorporated into oligonucleotide strands to improve the thermal stability and nuclease resistance of MBs for incubation at 37 ° C. The designed MBs were custom synthesized and purified by Sigma-Aldrich. An aqueous solution of MBs in PBS was vigorously mixed with a lipid formulation of dioleoyl-3-trimethylammonium propane (DOTAP), cholesterol, phosphatidylcholine (POPC), and 1, 2-distearoyl-sn-glycero-3-phosphoethanolamine-poly(ethylene glycol) (DSPE-PEG) in 200 proof ethanol, and then sonicated for 5 min using an ultrasonic bath. The MB/lipid mixture was subsequently injected into PBS, vortexed, and sonicated for 5 min. Finally, it was dialyzed with a 20 kDa MWCO dialysis bag to remove free MBs. After EVs were captured on magnetic beads, they were incubated with the prepared MBs for 2 h at 37 °C before imaging.

### Flow cytometry and fluorescence microscopy

tEVs from U251 GBM cells were stained with a lipophilic fluorescent dye, SP-DiOC18(3) (ThermoFisher Scientific) for 20 min, the excess dye was washed out. 100 μL fluorescent EVs (~10^11^ particles/mL) were then spiked into 500 μL of serum (~10^12^ particles/mL) and 100 mL of urine (~10^9^ particles/mL) from healthy donors. Each of the samples was processed by iSUF and recovered in 100 μL of PBS (stage 1 to 3). Non-spiked fluorescent tEVs were also captured on functionalized beads as a positive control. After washing with PBS, the captured EVs were analyzed by imaging flow cytometry (Amnis, ImageStream^X^ Mark II Imaging Flow Cytometer, Luminexcorp, Austin, TX); also images were taken using a fluorescence microscope (Nikon Eclipse Ti Inverted Microscope System) with a 100× oil immersion lens. For comparison, samples were also processed by UC and resuspended in 100 μL PBS. For total RNA quantification, captured EVs were lysed, and RNA was extracted and quantified using the same procedures mentioned above.

### Definition of terms used in this study

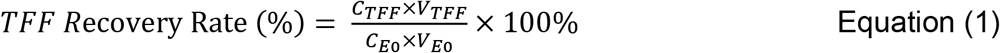

Where V_E0_ and C_E0_ are the initial volume and concentration of the sample; V_TFF_ (in 2 mL) and C_TFF_ (measured by qNano pore size [NP150, NP300, NP600, NP800, and NP1000]) are the final volume and concentration of the TFF product.

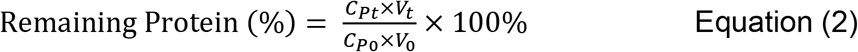

Where V_0_ and C_P0_ are the initial volume of the sample and concentration of free proteins in the sample, respectively; V_t_ and C_Pt_ are the volume of the TFF retentate and concentration of free proteins in the retentate at a specified time t, respectively. Concentration of free proteins was measured using the bicinchoninic acid (BCA) assay. The free protein removal efficiency can be calculated by subtracting the percentage of remaining protein in the final TFF product from 100%.

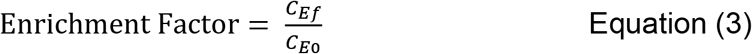

Where C_E0_ is the EV concentration in the initial sample; C_Ef_ is the EV concentration in the final iSUF product. Concentration of samples were measured by qNano pore size [NP150, NP300, NP600, NP800, and NP1000]).

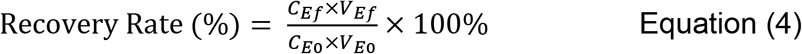

Where V_E0_ is the initial volume of the product; V_Ef_ is the volume of the final iSUF product.

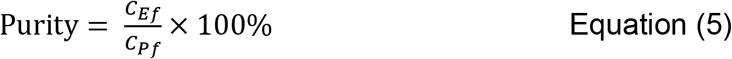

Where C_Ef_ is the final concentration of the iSUF product (in 100 μL); C_Pf_ is the remained free protein concentration in the product.

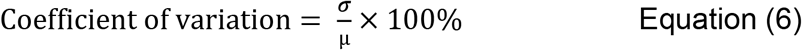

The coefficient of variation (CV) is a measure of relative variability. It is the ratio of the standard deviation (*σ*) to the mean (μ)

### Statistical Analysis

Data are expressed as the mean ± STD. A significant test between different mean values was evaluated using one-way ANOVA in JMP Pro 16 software provided by The Ohio State University. Differences between samples were considered statistically significant for p < 0.05.

## Results

### Optimization of the iSUF platform

To overcome current limitations of the enrichment and purification of EVs and on-demand EV subpopulation characterization, we developed the iSUF platform (**Fig. 1 A**) which includes three stages: (1) tangential flow filtration (TFF) for the enrichment and purification of EVs, (2) centrifugation for further enrichment of EVs, and (3) immunomagnetic affinity selection for desired EV subpopulation isolation. We used the iSUF platform to process various volumes (0.5-100 mL) of different biofluids (CCM, serum, and urine).

To design stage 1, many parameters of TFF processing required optimization, including the selection of membrane pore size (MWCO), sample processing temperature, sample flow rate, pressure, and sample protein concentration. We tested membrane filters with two MWCO sizes (300 and 500 kDa) to determine the optimal MWCO that maximizes the removal of free proteins and nucleic acids while reducing processing time. Our experiments showed that 500 kDa filter membranes were able to remove up to 99 % of free proteins with over 99% EV recovery rate (Equation 1, **Supplementary Fig. 1**). When a 300 kDa membrane filter was used, only 80 % of free proteins were removed. Moreover, a 500 kDa membrane filter was chosen since it processed samples 2~3 times faster than a 300 kDa membrane filter (**Supplementary Table 1**). The TFF stage 1 of iSUF was run at 4°C to minimize EV degradation^39^. We further tried to optimize sample processing time, which was highly dependent on the flow rate. The flow rate was linearly associated with the shear rate generated by the filter based on the manufacturer’s protocol (Repligen), which exerted a shear force on the EVs. Then, we used a flow rate of 35 mL/min to maintain a shear rate below 5000/s^40^. High flow rates increased the system pressure, mainly when the protein concentration of the sample was high (> 15 mg/mL). We kept the pressure of the system below 10 psig to avoid leakage and maximize the lifespan of the 500 kDa filter. We used dilutions of fetal bovine serum (FBS) to test the effect of protein concentration on system pressure at 35 mL/min. Our results showed that protein concentration must be equal or lower than 15 mg/mL to maintain the pressure of the system below 10 psig to protect the filter (**Supplementary Fig. 2**).

At stage 2 of iSUF, EVs were loaded into centrifugal filter units and were spun down at 3,000 x *g*. We compared the recovery rates of EVs and processing time for different filter pore sizes. A 3 kDa filter unit obtained over 99% recovery rate and took 60 min to spin down, while a 10 kDa unit obtained a 95% recovery rate in 20 min, and a 30 kDa filter obtained only a 70% recovery rate and took 15 min to spin down (**Supplementary Fig. 3**). We selected the 10 kDa filter unit to maintain a high recovery rate while reducing sample processing time. To enrich tEVs (stage 3 of iSUF), we tested incubation of 3 μm magnetic beads with different concentrations of biotinylated antibodies (10, 20, 100 μg/mL) for 1 h and 2 h at RT, and overnight at 4 °C. Overnight incubation with 100 μg/mL of antibody exceeded the bead-antibody binding efficiency, while 20 μg/mL of antibody was able to yield a high bead-antibody binding efficiency (>90%). Then, we examined the volume ratio of beads to EVs at 5 μL /100 μL, 20 μL /100 μL, and 80 μL /100 μL during the bead/tEVs incubation step. The 20/100 μL ratio achieved the highest bead-tEVs capture efficiency (>90%).

### iSUF platform for processing biofluids

A flow rate of 35 mL/min for stage 1 of iSUF was applied since the protein concentrations of the different biofluids were below 15 mg/mL (**Supplementary Table 2**). We tested the ability of our platform to purify and enrich EVs from three different biofluids (i.e., CCM, serum, and urine; **Fig. 1B**). 50 mL of CCM, and 100 mL of urine were enriched to 7 mL. Subsequently, PBS diafiltration buffer was used to remove the remaining protein contaminants from CCM and urine in 80 min and 100 min, respectively. The percentage of remaining protein is defined as the mass of free proteins in the TFF retentate at a specified time divided by their initial mass in the sample (Equation 2). For serum, the initial high concentration of proteins (> 80 mg/mL) required an initial dilution of 0.5 mL of the sample in 7 mL of PBS. Subsequently, PBS diafiltration buffer was used to remove the remaining protein contaminants from serum in 120 min. We first tested iSUF with a 10% BSA solution for which an SDS-PAGE gel showed extensive removal of albumin (**Supplementary Fig. 4**). Moreover, analysis of the purified samples by an SDS-PAGE gel indicated that iSUF removed BSA from CCM, human serum albumin (HSA) and globulins from serum, and Tamm-Horsfall glycoprotein (THF) and HSA from urine. (**Supplementary Fig. 5, 6, 7**).

### Concentration, size distribution and microscopy characterization of EVs

For 50 mL of CCM, 0.5 mL of serum, and 100 mL of urine, the enrichment factors were 489 ± 18, 4.8 ± 0.1, and 942 ± 19, respectively (n = 5; **Fig. 2A**) (Equation 3). Accordingly, the EV recovery rate (the ratio of the total number of EVs post-processing by iSUF to the total number of EVs pre-processing) was 98% ± 3.6%, 96% ± 2.0% and 94% ± 1.9% for CCM, serum, and urine, respectively (Equation 4). Considering that EVs are heterogeneous in size, we also tested the EF across a wide size range of EVs (70 nm - 1μm) in CCM with similar results (n = 5; p > 0.05; **Fig. 2B**). Moreover, the EV concentrations obtained by iSUF was greater than the concentrations obtained by UC. This difference was consistent across the 70 nm to 1 μm size range (n = 5; p < 0.05; **Fig. 2C**), with iSUF enriching EVs at two to three orders of magnitude higher than UC (approximately 10^11^ EVs concentrated by iSUF and 10^9^ for UC). Similar results were obtained with comparisons for the enrichment of EVs present in serum and urine. With iSUF, EVs from serum and urine were enriched almost at the same level (10^12^ EVs/mL; **Fig. 2D**). We confirmed the presence of EVs by using atomic force and electron microscopy on the different biofluids processed. The majority of isolated EVs exhibited a round morphology with heterogeneous size distribution. Cryo transmission electron microscopy images (TEM) images of isolated EVs showed the presence of a double-layered lipid membrane, a representative characteristic of EVs. (**Fig. 2E & 2F**).

**Figure 2.**
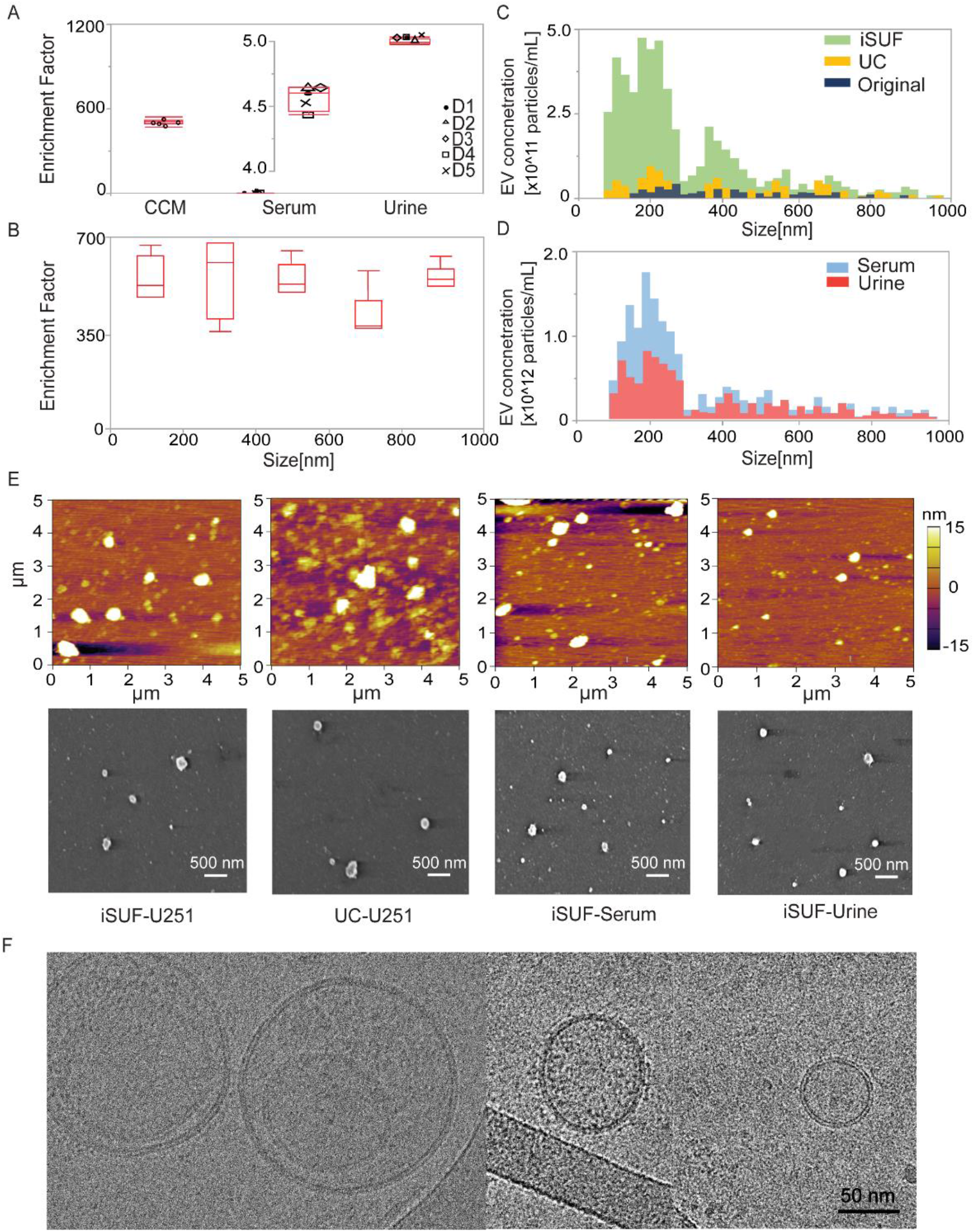
iSUF enrichment performance and characterization of EVs separated from CCM, serum and urine using iSUF. A) Enrichment factor (EF) for EVs present in CCM, serum, and urine after iSUF (n = 5 for each biofluid; p>0.05) (mean ± STD). Enrichment factors (EFs) were calculated as the ratio of EV concentration in biofluids present before and after iSUF processing. B) EFs for different size ranges of EVs for CCM (n = 3; p > 0.05) (mean ± STD). C) qNano measurements of the size distribution and concentration of EVs in the original CCM, after iSUF processing, and after UC processing. The EV concentration after iSUF was significantly higher than in the original CCM and after UC processing (n = 3; p < 0.05) (mean ± STD). D) qNano measurements of the size distribution and concentration of EVs in serum and urine after iSUF processing. E) AFM and SEM images of EVs from U251 CCM after iSUF processing and UC processing. Images were also obtained for EVs in serum and urine after iSUF processing. F) Transmission-EM images of EVs present in CCM after iSUF processing.

To further test our iSUF platform, we performed comparative studies with different commercially available EV isolation methods: qEV, TEI, and UC. 0.5 mL of serum from healthy donors were processed with different EV isolation platforms. EVs demonstrated similar smaller and larger size distribution for all platforms, but iSUF obtained a higher EV concentration than the other methods within the 70 nm - 1 μm size range (**Fig. 3A**). We also compared the mean size of subpopulations of EVs based on their physical characteristics using different methods^38^. **Fig. 3B** shows the mean size of small EVs (sEVs) and medium/large EVs (m/l EVs). For both size ranges, EVs processed by iSUF were significantly smaller than UC (n = 5; p < 0.05). For l/m EVs, EVs purified by UC were larger than all other techniques (n = 5; p < 0.05). Then, we compared the total concentration and purity of isolated EVs using different methods. iSUF enriched EVs significantly more efficiently than all the tested methods (n = 5; p < 0.05; **Fig. 3C**). The concentration of EVs enriched by iSUF was 51 ± 18, 7.0 ± 2.0, and 56 ± 21 times higher than qEV, TEI, and UC. The purity of isolated EVs was normalized and evaluated in terms of the ratio between EV concentration and remaining contaminating protein concentration present in samples after purification using the different methods (Equation 5). The purity of the isolated EVs by iSUF was 100-10000 times higher than the different tested methods (**Fig. 3C**).

**Figure 3.**
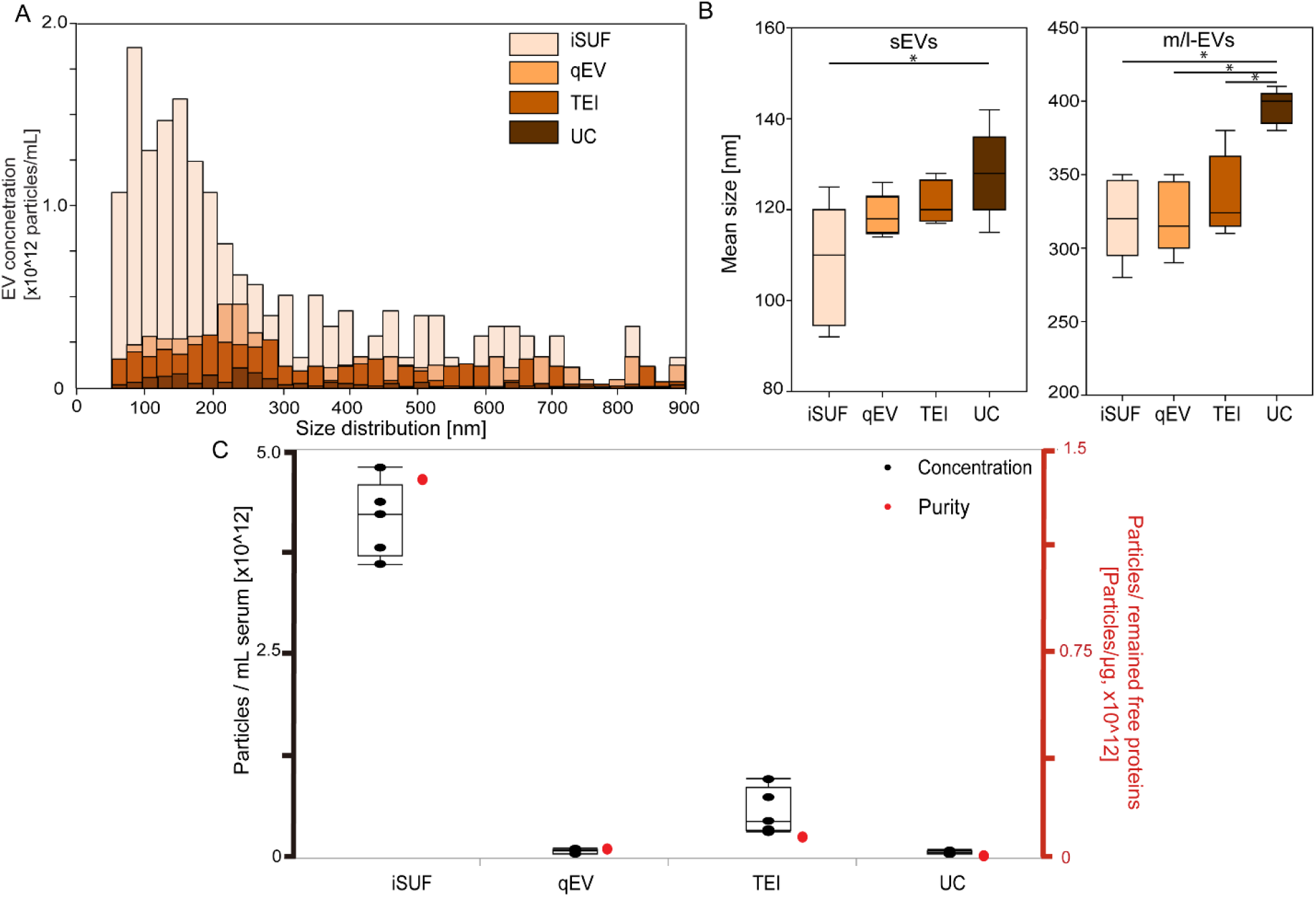
Comparison of size distribution and purity of EVs isolated with different platforms. A) Size distribution of EVs isolated from serum using different platforms. iSUF demonstrated the highest EV concentration within the 70 nm – 1 μm size range. B) Left. Mean size boxplot for small EVs (sEVs). sEVs isolated using qEV and TEI demonstrated no differences with iSUF or UC, UC-sEVs showed larger mean size than iSUF-sEVs (n = 5; p < 0.05) (mean ± STD). Right. Mean size boxplot for medium/large EVs (m/l EVs). UC-m/l EVs were larger than iSUF, qEV and TEI m/l EVs (n = 5; p < 0.05) (mean ± STD). C) Concentration and purity of isolated EVs using different platforms. Black boxplots were the absolute concentration of EVs isolated from 0.5 mL of serum (left-axis); red dots corresponding to the right y-axis were purities defined as the EV concentration divided by the remaining free protein concentration. iSUF isolated EVs from serum most efficiently with high purity (n = 5; p < 0.05) (mean ± STD), qEV yielded pure EVs, but at a lower concentration, TEI isolated more EVs but at relatively low purity, and UC recovered EVs with the lowest concentration and purity.

Repeatability experiments for the EV recovery rate and protein removal efficiency from samples processed by iSUF were also conducted (CCM, n = 10; serum, n = 10; Urine, n = 10). Both sEVs and m/l EVs in all biofluids demonstrated a mean value above 94% for the EV recovery rate. The recovery rate of sEVs in CCM, serum, and urine exhibited 4.2%, 3.7%, and 2.4% coefficient variations, respectively, while m/l EVs showed 3.7%, 3.3%, and 2.5% coefficient variations (Equation 6). Protein removal efficiencies were also tested, CCM, serum, and urine presented up to 99% removal efficiency with coefficient variations of 0.9%, 4.0%, and 2.1% (**Supplementary Fig. 8**).

### Molecular content quantification and characterization of EVs

We quantified the total amount of protein and RNA present in EVs isolated by iSUF for different biofluids. For CCM, the quantity of RNA obtained was 11 ± 8.2 ng/mL, thus giving a 9-fold higher concentration of RNA when compared to UC (**Fig. 4A**). For protein analysis, the protein concentration was 6 ±1.8 μg/mL, almost 8-fold more protein obtained than when the same biofluid was processed by UC (**Fig. 4B**). Total RNA and protein quantification were carried out for serum and urine from 5 healthy donors. 0.5 mL of serum and 100 mL of urine processed by iSUF produced 54 ± 40 ng/mL and 0.3 ± 0.1 ng/mL of RNA, respectively (**Fig. 4C**). The protein concentration was 1250 ± 480 μg/mL for serum and 3.1 ± 3.0 μg/mL for urine (**Fig. 4D**). The differences in protein concentration of EVs in original CCM, processed by UC, and processed by iSUF were also demonstrated using CD63 and CD9 western blot analysis. Also, the expression level of CD9 and CD63 in original urine and serum samples, and after iSUF processing demonstrated superiority enrichment and purification performance of the iSUF (**Fig. 4E**). Full-length western blot images are displayed in (**Supplementary Fig. 9**). Moreover, we compared the RNA content obtained from EVs enriched from 0.5 mL of serum using different commercially available methods (**Supplementary Table 3**). The total RNA content obtained using iSUF was 54 ± 41 ng, which was significantly higher than other methods that only obtained 10 ± 9.8 ng (**Fig. 4F**).

**Figure 4.**
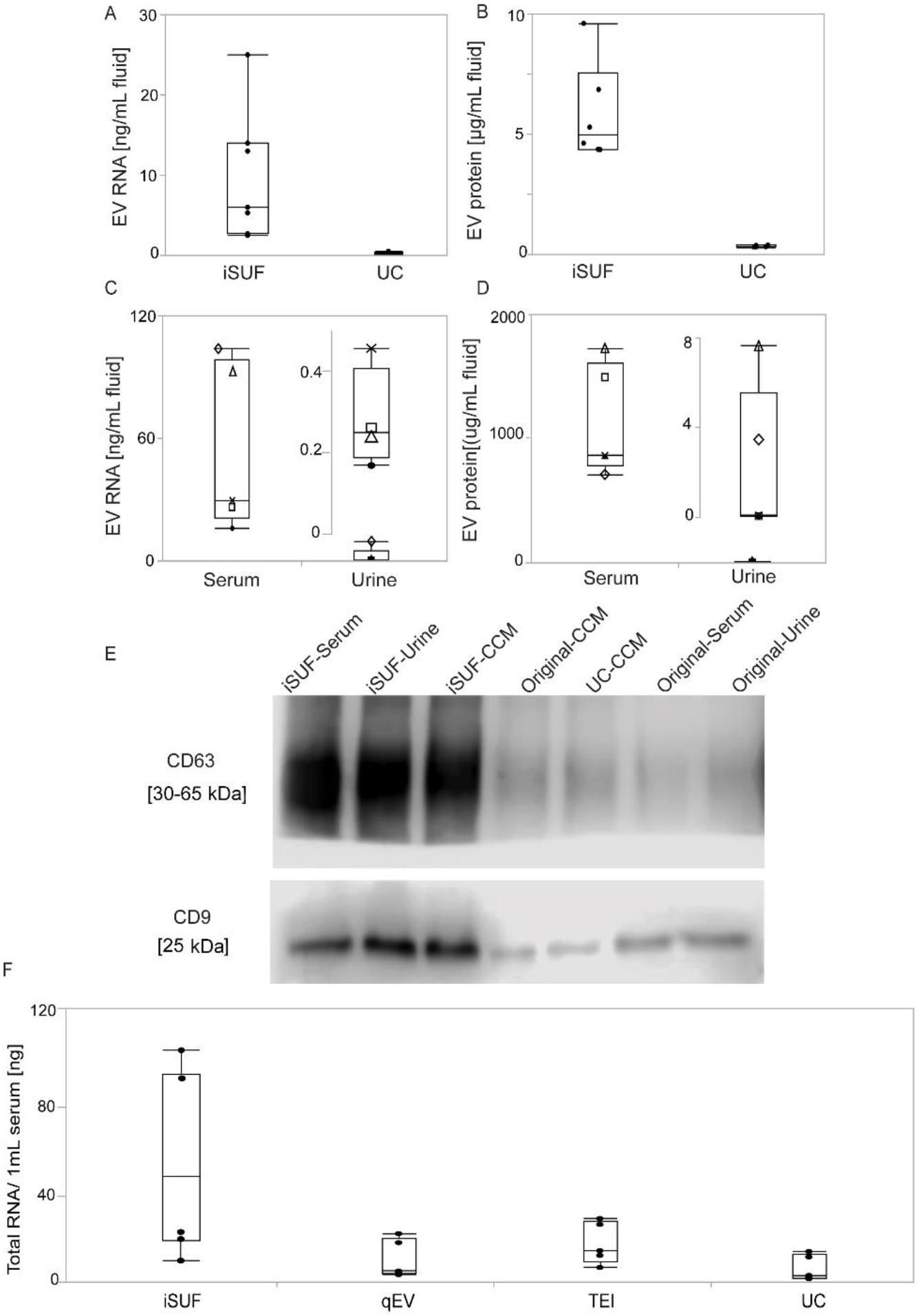
Characterization of protein and RNA content of EVs isolated from different biofluids. A) Quantity of total RNA extracted from CCM-EVs using UC and iSUF (n = 6; p < 0.05). B) Comparison of proteins extracted from CCM-EVs using UC and iSUF (n = 6; p < 0.05). C) Quantification of total RNA extracted from serum and urine EVs isolated by iSUF. D) Quantification of EV proteins isolated from serum and urine EVs isolated by iSUF. E) Western blot analysis of CD63 and CD9 expression in original CCM, CCM-EVs processed using iSUF and UC, original serum and urine, serum and urine-EVs processed using iSUF. The expression level of CD63 and CD9 was calculated using an equal mass (40 μg) of protein lysates from samples. F). Total RNA of EVs isolated using different EV isolation methods. Black boxplots were the absolute amount of RNA isolated from 0.5 mL of serum. iSUF isolated the most RNA contents from serum EVs (n = 5; p < 0.05) (mean ± STD).

Next, we processed different volumes of serum and urine with iSUF to identify the equivalent volumes of biofluid that will produce comparable concentrations of EVs, total RNA, and total protein. We started with 0.5 mL of serum and different volumes of urine (i.e., 100, 75, 65, 10 mL). For different volumes of urine and 0.5 mL of serum, the enriched concentrations of EVs were comparable (~ 10^12^ particles/mL, **Supplementary Fig. 10**). The total RNA content obtained from EVs from serum and urine also exhibited comparable values of 60 ± 35 ng and 75 ± 40 ng, respectively. However, analysis of protein content of the isolated EVs from both biofluids showed that the protein content of EVs isolated from urine was 10-fold lower than the protein content of EVs in serum, which was 0.4 ± 0.1 mg and 40 ± 10 mg, respectively.

### Immunomagnetic affinity selection of the iSUF platform

tEVs are surrounded by many normal EVs (nEVs) that require removal to perform accurate molecular analysis^41^. One way to isolate tEVs is to exploit the presence of specific surface markers^42^. Our iSUF platform enables the separation of subpopulations of EVs by capturing them on magnetic beads through immunoaffinity. To demonstrate our approach, we used a model system that consisted of spiking fluorescently labeled tEVs from a cancer cell line (i.e., U251 EVs) in 0.5 mL of serum and 100 mL of urine from healthy donors (n = 3). 3-μm magnetic beads were functionalized with EGFR as the capture antibody. All samples were processed through all stages of iSUF, and the captured subpopulation of EVs was characterized and quantified using TIRF and imaging flow cytometry (**Fig. 5A**). A comparable fluorescent signal was obtained between the positive control (tEVs directly from U251 cell supernatant) and spiked samples processed in serum and urine, which verified the high recovery efficiency of tEVs (88 ± 1 for serum, and 93 ± 2 for urine), and their purity. The biofluids spiked with tEVs were also processed by UC, which yielded significantly lower fluorescent signal (3.1 ± 0.1 times lower, **Fig. 5A**). Furthermore, to determine tEV RNA isolation efficiency by iSUF, lysed U251-EVs (positive control) and bead-isolated U251-EVs were extracted. iSUF showed over 90% RNA isolation efficiency for the EGFR^+^ tEVs (**Fig. 5B**). Moreover, the RNA content of bead-extracted tEVs from serum using the iSUF platform was 7-fold higher than UC, which was mainly attributed to the high yield of total EVs recovered in iSUF (**Fig. 5C**). The high RNA content of tEVs from serum isolated using iSUF also confirmed the ability of iSUF to enrich and retrieve tumor targets from mixed populations of EVs.

**Figure 5.**
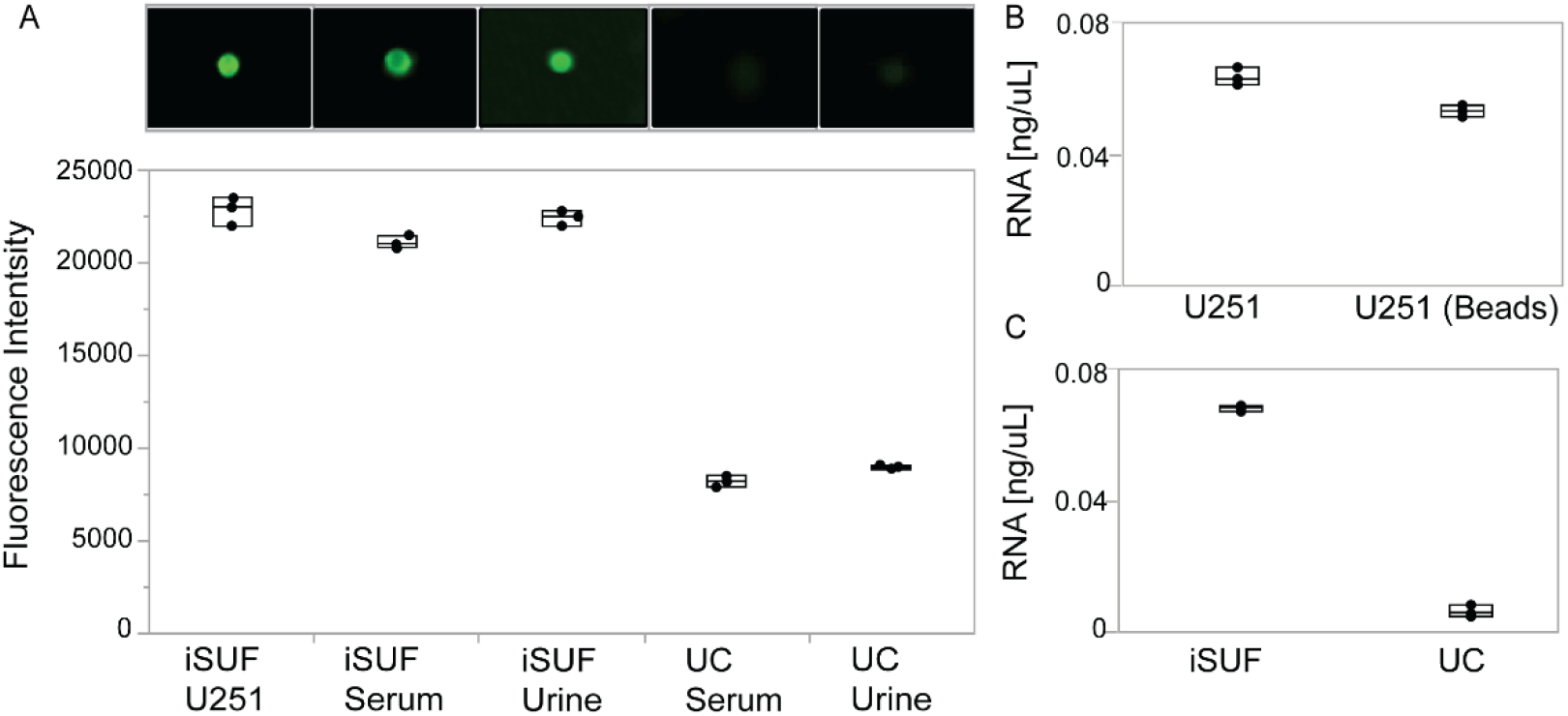
Characterization of stained tEVs isolated from U251-CCM and spiked in different biofluids. A) Upper. Fluorescence images of tEVs on magnetic beads using TIRF microscopy. Lower. Comparison of fluorescence intensities quantified using imaging flow cytometry. Stained U251-EVs isolated by iSUF was a positive control. tEVs isolated from serum and urine samples using iSUF demonstrated no significant difference (n = 3; p > 0.05), while UC isolated tEVs differed with iSUF and the positive control (n = 3; p < 0.05) (mean ± STD). B) Quantification of total RNA isolated from U251-EVs before and after bead extraction showed no differences (n = 3; p > 0.05). C) Total RNA extracted from serum samples using iSUF was significantly higher than using UC (n = 3; p < 0.05) (mean ± STD).

### Detection of proteins and miRNA in EVs from clinical samples

The emergence of targeted therapies require a precise characterization of the molecular subtypes of a patient’s tumor^43^. For BC patients (e.g., luminal A/B, triple-negative), the molecular subtype is typically determined by testing a tissue biopsy for the presence or absence of three essential proteins: estrogen receptor (ER), progesterone receptor (PR), and human epidermal growth factor receptor 2 (HER2)^44^. However, a tissue-based tumor profile is subject to sampling bias, provides only a snapshot of tumor heterogeneity, and cannot be obtained repeatedly^45^. These limitations restrict early cancer detection and significantly contribute to the overdiagnosis and overtreatment of BC patients^46^. The analysis of EVs present in biofluids of cancer patients provides a minimally invasive method to quantify different biomarkers that would enable precise diagnosis or response to treatment^47^. To test the potential use of the iSUF platform in screening biomarkers for BC, we processed 0.5 mL of serum from 10 metastatic BC patients and quantified the expression levels of HER2, CD24, and miR21 on patient isolated EVs^48–50^. After stage 3 of iSUF, magnetic beads were isolated using a magnet and incubated with detection antibodies and MBs. Then, the magnetic beads were processed with imaging flow cytometry and TIRF microscopy. Similar volumes of serum from healthy donors were processed and analyzed. The age of the healthy donors was between 21 and 68 years old. Although the age range was wider than from our BC patient cohort, a comparison of levels of expression for HER2, CD24, and miR21 between healthy donors samples did not show statistically significant differences (p > 0.05, **Supplementary Fig. 11**). We obtained representative images and histograms that quantify the number of beads and their corresponding fluorescent intensities (**Fig. 6A** & **6B**) for BC patient and healthy donor samples. Both protein and miRNA show significantly higher expression in BC patient samples. (p < 0.05; **Fig. 6B**), which suggested that iSUF can differentiate tumor biomarkers (**Supplementary Table 4**). Moreover, we have characterized the CD9 and CD63 levels of protein expression in serum samples from both cohort of samples. Although in this case, similar levels of expression were obtained (p > 0.05, **Fig. 6A & 6B**).

**Figure 6.**
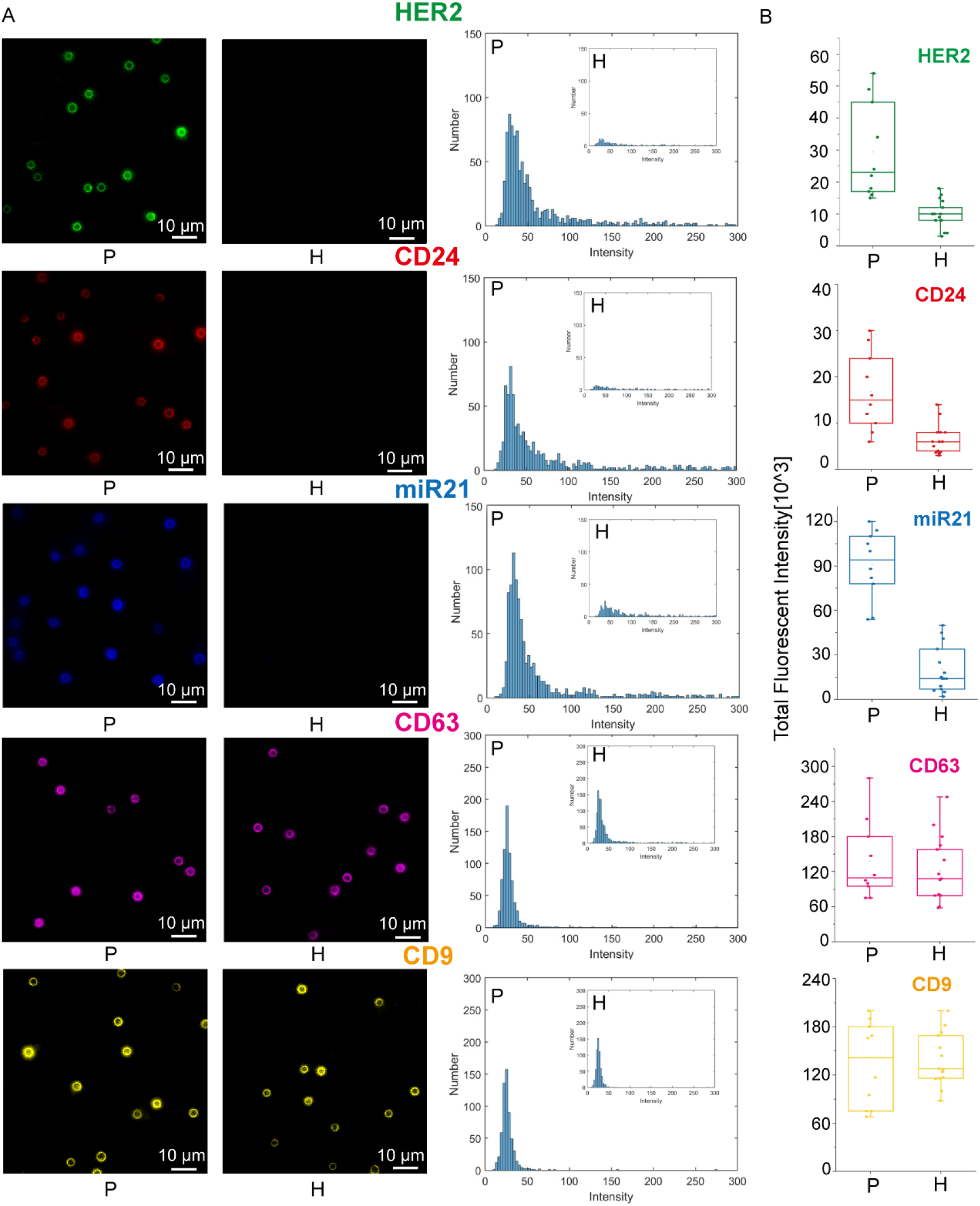
Characterization of tEVs isolated breast cancer (BC) patient serum using iSUF. A) Left. Characteristic fluorescence images of HER2, CD24, miR21, CD63, and CD9 expression on isolated EVs immobilized on magnetic beads from BC patients and healthy donors. Right. Fluorescence intensity histograms of HER2, CD24, miR21, CD63, and CD9 of isolated EVs for BC patients (n = 10) and healthy donors (n = 14). B) Total fluorescence intensity quantification of HER2, CD24, miR21, CD63, and CD9 expression level on isolated EVs from BC patients and healthy donors. Patients demonstrated higher expression levels of HER2, CD24, and miR21 than healthy donors (p < 0.05). Patients and HDs demonstrated similar expression level of CD63 and CD9 (p > 0.05). P: patient; H: healthy donor.

## Discussion

Recently, EVs have been explored for diagnostic and therapeutic applications, including liquid biopsy assays for cancer diagnostics, and nanocarriers for drugs and nucleic acids^51,52^. The large number of tEVs compared to other rare biomarkers (i.e., circulating tumor cells, CTCs) makes them more statistical reliable^53^. Innovative methods developed for the enrichment and purification of EVs should remove all contaminants (e.g., free proteins), have a high yield, work amongst different biofluids, and maintain the integrity of the EVs. We have engineered a new platform that includes a TFF enrichment and purification stage, a centrifugal-unit enrichment stage, and a magnetic separation stage for specific isolation of subpopulations of EVs (e.g., tEVs). In stages 1 and 2, iSUF performs enrichment and purification of all EVs (e.g., tEVs and nEVs). In stage 3, tEVs are isolated based on their tumor-specific surface markers. 300 kDa and 500 kDa TFF filters were initially selected for this study because the molecular weights of the major free proteins in CCM (e.g., BSA, 65 kDa), serum (e.g., HSA, 65 kDa) and urine (e.g., THP, 98 kDa) were below 300 kDa^54–56^. After testing, a 500 kDa TFF filter was finally chosen for subsequent experiments because of its minimal EV loss and faster processing time (~ 2 h).

Different biofluids required different TFF diafiltration buffer volumes and different scale of TFF hollow fiber filters to remove most contaminants. CCM and urine contain relatively low concentrations of protein contaminants^57,58^, so a small total diafiltration buffer volume was required to remove the majority of free proteins. Unlike CCM and urine, the serum has a significant amount of free proteins^59^. Therefore, a large total diafiltration buffer was necessary to remove the free protein in serum. iSUF was able to achieve up to 99 % of free protein removal efficiency. Free proteins removed by iSUF were shown as decreased bands on the SDS-PAGE gel. Other bands were increasing on the SDS-PAGE gel because those correspond to different proteins present on EVs, which were enriched by the iSUF^60–64^. Also, there are Very-low-density lipoproteins (VLDL)) that overlap in size with EVs that also might be enriched^2,65,66^.

Although TFF can concentrate and purify samples, the final product volume is mostly dependent on the dead volume of the specific TFF system. Therefore, further enrichment is necessary to concentrate the samples (e.g., spin down to - 100 μL) for applications such as the diagnosis of rare biomarkers^67^. Centrifugal units with different MWCOs (3, 10, and 30 kDa) were evaluated in terms of processing time and yield in our study. A larger MWCO size required shorter centrifugation time, but more EVs were lost because prolonged high-speed centrifugation elongated EVs into an oval shape which made them squeeze through the filter membrane^68,69^. The centrifugal unit with the 10 kDa MWCO was optimal with the shortest processing time and highest yield.

We also compared EVs purified from serum and urine samples of healthy donors. EVs from urine and serum can serve as prognosis biomarkers for clinical analysis^70,71^. Serum is the most commonly used biofluid in the clinical setting with the highest EV concentration^72^. Compared to serum, urine collection is minimally invasive and can be obtained in larger volumes^73^, but it usually suffers from much lower EV concentration^74^. Interestingly, using our iSUF platform, the EV RNA concentration in the final products of urine samples (originating from ~ 100 mL of collected sample volume) were comparable to those of serum samples (originating from 0.5 mL of collected sample volume). This suggests that clinical diagnosis by urinary EVs is possible. However, there is still a concern towards urinary EV collection because of large variabilities in the urine volume and its EV concentration. More efforts are needed to come up with a gold standard protocol for urine collection and processing. For protein in urinary EVs, we found that the amount of protein was lower than the amount of protein obtained from serum EVs, which might be explained by the degradation of EV membrane proteins by urine proteases^75^.

In this study, we compared the performance of iSUF with three different EV isolation techniques (TEI, qEV, and UC) to determine EV concentration, purity, and quantity of RNA recovered. As other authors have discussed^76,77^, we found that UC, a traditional method for EV isolation, has raised concerns about the integrity, yield, and purity of EVs after purification. Interestingly, we found UC-isolated EVs were larger than other platforms, one possible explanation is the presence of extensive levels of protein aggregates that cause bias. For our UC method, washing steps using multi-rounds ultracentrifugation were not applied, since it was shown that purity of the samples did not increase significantly when adding washing steps, and a significant loss of enriched EVs could occur^54,78–82^. Moreover, although qEV obtained a relatively pure product, its recovery rate was low; TEI obtained a relatively higher EV number, while retaining massive amounts of free protein (**Fig. 3C**). Therefore, low EV concentration and purity will impact the accuracy of molecular analysis of EVs^83^.

We used the MISEV 2018 guidelines to define EVs: small EVs (sEVs) and medium and large EVs (m/l EVs)^38^. Following these definitions, we found that the concentration and average size of EVs in patient samples and healthy controls varied (**Supplementary Table 4**). Different factors may explain these differences. Subtypes and stages of BC in the patient cohort, may contribute to the difference in the metrics. Moreover, we obtained our patient samples from a biobank, we had minimal control of the storage conditions, thus it was possible that some degradation of the samples could occur. It is known that factors such as material of the container, temperature of storage, and cycles of freezing and thawing may also influence the concentration and mean size of EVs^84–87^. Considering all these aspects, the size distributions and concentrations of EVs that we obtained for the different samples are within the reported values in the literature. Previously concentrations of EVs in serum from cancer patients are between 10^9^ – 10^11^ particles/mL ^88–91^.

Differentiating EV particles and other non-EV particles (e.g., lipoproteins) when measuring size distribution and concentration is challenging for all current methods, including TRPS and nanoparticle tracking analysis NTA^65,92–96^. This is because the size range of different lipoproteins (e.g., 7 – 13 nm (HDL); 21 – 27 nm (LDL); and 30 – 90 nm (VLDL))^97^ may overlap with the reported size ranges for sEVs and m/l EVs^38^. Thus, current methods for EV size characterization and concentration have certain limitations^87,98–100^. For iSUF, we used the TRPS method to measure size distribution and concentration of EVs. Different nanopore stretchable membranes (NP150, NP300, NP600, NP800, and NP1000) were selected, covering a broad size range of EVs such as 70 nm to 1000 nm^101^. The 70 nm cut-off were chosen since EVs in samples processed by iSUF, exhibited a range above 70 nm. Choosing a small stretchable membrane such as NP100 and NP80 that have limits of detection of 50 nm and 40 nm, respectively, did not change the size ranges of EVs in our samples after iSUF (**Supplementary Fig. 12**). Thus, iSUF intrinsically filters out the majority of lipoproteins and EVs that overlap in size. This effect is beneficial since we are trying to characterize EVs with minimal contribution of lipoproteins. We consider the 70 nm cut-off used for the TRPS measurements as appropriate. Moreover, above this value the different characterization metrics are considered accurate. The particle count measurements can verify this for each one of the methods used in our study (**Supplementary Fig. 13**). A linear relationship indicates an accurate measurement^30^. In **Fig 3**, we found that the lower limit of detection varies for the different methods, indicating an intrinsic loss of EVs during processing samples with different methods. However, iSUF shows better recovery for sEVs and m/l EVs. The wide upper range showed by UC could be indicative that we measure aggregates of EVs^102–105^ Our measurements are in agreement with MISEV 2018 since sEVs, and m/l EVs are in the 100 ~ 150 nm and larger than 200 nm, respectively^38,62,106–110^.

We are interested in the enrichment and isolation of tEVs to characterize tumor-related proteins and RNAs. Like other immunoaffinity methods^111–113^, iSUF captured and isolated EVs using specific tumor surface proteins. iSUF differentiated metastatic BC patients from healthy donors by detecting significantly higher expression levels of proteins and RNA biomarkers present in EVs (e.g., HER2, CD24, and miR21). Based on previous reports, HER2 and miR21 are cancer-associated protein and microRNA species, and are known to be overexpressed in metastatic BCs^114,115^. Compared to miR21 and HER2, CD24 is relatively less investigated in BC but was previously identified as being released from BC stem cells^116^. Furthermore, a recent study indicated that serum CD24 is elevated among BC patients^117^. Moreover, it is important to note that one of these biomarkers may not be a reliable predictor of BC alone. However, the combination of several biomarkers can serve as a tool for BC risk assessment.

In conclusion, iSUF was proposed for rapid, efficient, and specific isolation of EVs from different biofluids. The EV recovery rate mean value was above 94%, with 90% specific isolation of tEVs and negligible concentrations of free proteins and nucleic acids. Although iSUF does not process a sample in the shortest time (80 min ~ 120 min), its versatility working with different biofluids, sample volumes, and high purity after sample processing constitutes unmatched advantages over current methods used in the field. Overall, we found that the iSUF platform isolated and enriched EVs from a scaled-up sample volume with high purity and yield in a sterile and quick manner simultaneously, and isolated tEVs with high specificity, while other current methods could not guarantee all of those conditions at the same time. Furthermore, we recognize that the iSUF platform has potentially broad clinical applications beyond liquid biopsies for cancer diagnosis or monitoring.

## Acknowledgments

We acknowledge all the patients and healthy volunteers who participated in this study. This work was supported by the U.S. National Institutes of Health (NIH) grants UG3TR002884 (ER). and R01HL126945, R01HL138116, and R01EB021926 (AFP). Additional support for E.R. was provided by the William G. Lowrie Department of Chemical and Biomolecular Engineering and the Comprehensive Cancer Center at The Ohio State University.

## Contributions

E.R and A.F.P. conceived the idea. E.R., L.T.H.N, and J.Z design the study. J.Z. performed most of the experiments and data analysis, prepared all tables and figures, and drafted the original manuscript with inputs from all authors. L.T.H.N established protocols for protein/RNA analysis and immunomagnetic beads and performed some of the experiments. R. H. performed equipment installation and calibration. N.W. performed sample collection and supported with editing the manuscript. X.W. developed the quantification code for the third stage of iSUF. K.K. designed molecular beacons for targeting miR21. A.F.P. supervised the TFF experiments and edited the manuscript. L.J.L. supervised the design of molecular beacons and edited the manuscript. E.R. supervised the whole study and edited the manuscript. All authors provided critical feedback and helped to shape the research, analysis, and manuscript.

## Supplementary Information

### Size distribution and concentration of EVs

A tunable resistive pulse sensing (TRPS) method (Izon Sciences, Boston, MA) was employed to quantify the size and concentration of EVs. 45 μL of biofluid was pipetted into nanopore membranes (NP150, NP300, NP600, NP800, and NP1000), and then pressure (10 mbar) and voltage (0.38V, 0.32V, 0.26V, 0.18V and 0.12V) were applied. Every single EV causes a resistive pulse that can be used to calculate EV size and concentration. Polystyrene nanoparticles of different known sizes and concentrations were used for calibration.

### Atomic force microscopy (AFM)

A clean mica substrate was vapor-phase coated with 3-aminopropyltriethoxysilane (APTES, Millipore Sigma, Burlington, MA) in a vacuum chamber and then dried overnight at 65 °C. Subsequently, 10 μL of purified EVs were incubated on the surface for 30 min at RT. Unbound EVs were extensively rinsed with PBS and then with DI water. The samples were air-dried again before imaging using an AFM (Asylum Research MFP-3D-BIO AFM, Oxford Instruments, Abingdon, United Kingdom).

### Scanning electron microscopy (SEM)

Clean coverslips were soaked in 0.25 mg/mL Zetag solution (BASF, Southfield, MI, USA) for 30 min, followed by overnight air drying at RT. Purified EVs were attached to the coated coverslip for 30 min at RT by physisorption. EVs were fixed in 2% glutaraldehyde (MilliporeSigma, Burlington, MA) and 0.1 M sodium cacodylate solution (Electron Microscopy Sciences, Hatfield, PA, USA) for 3h. After washing with 0.1 M sodium cacodylate solution, EVs were incubated in 1% osmium tetraoxide (Electron Microscopy Sciences) and 0.1 M sodium cacodylate for 2h. The sample was subsequently rinsed with 0.1 M sodium cacodylate solution before dehydration in increasing concentrations of ethanol (50, 70, 85, 95, and 100%, ThermoFisher Scientific) for 30 min each. Next, the samples were transferred to a CO_2_ critical point dryer (tousimis, Rockville, MD, USA). Finally, the samples were coated with ~ 2 nm of gold using a sputtering machine (Leica EM ACE 600, Buffalo Grove, IL) and imaged using SEM (Apreo ii, FEI, Thermo Fisher Scientific).

### Transmission electron microscopy (TEM)

3 μL EVs purified from CCM using iSUF were applied to a glow discharged lacey carbon-coated copper grid (400 mesh, Pacific Grid-Tech, San Francisco, CA) and flash-frozen in liquid ethane using an automated vitrification device (FEI Vitrobot Mark IV, FEI, Hillsboro, OR). The sample was then visualized in a Glacios™ Cryo-TEM (ThermoFisher Scientific).

### Protein extraction and quantification

EV samples were lysed in RIPA buffer (Abcam) with the addition of Thermo Scientific™ Halt™ Protease and Phosphatase Inhibitor Cocktails for 15 min on ice. EV samples (with/without lysis) were then pipetted to a 96-well plate, and their protein concentrations were quantified using a Pierce™ Rapid Gold BCA Protein Assay kit (ThermoFisher Scientific). EV protein concentration was determined by subtracting the amount of free protein (without lysis) from the total (with lysis) in the purified EV sample.

### Sodium dodecyl sulfate-polyacrylamide gel electrophoresis (SDS-PAGE)

Proteins in the final product were denatured and reduced in the presence of NuPAGE Reducing Agent in NuPAGE LDS Sample Buffer at 95 °C for 10 min. The proteins were then separated in a mini gel tank (ThermoFisher Scientific) using NuPAGE 4-12% Bis-Tris Protein Gel in NuPAGE MOPS SDS Running Buffer for 50 min at 200 V. After separation, the proteins were stained with Coomassie Brilliant Blue G-250 Dye.

### Western blotting

After separation by SDS-PAGE, proteins were transferred onto a polyvinylidene fluoride (PVDF) membrane (ThermoFisher Scientific) and then blocked with 3% bovine serum albumin (BSA) and 0.05% Tween in PBS for 1 hr at RT. Primary antibodies against tetraspanin surface markers such as CD63, CD81, and CD9 were incubated with the EVs overnight at 4 °C (Santa Cruz Biotechnology, Inc Dallas, TX, USA). The next day, the PVDF membrane was incubated with an HRP conjugated secondary antibody for 1h at RT. Finally, the sample was incubated with SuperSignal™ West Femto Maximum Sensitivity Substrate for 5 min at RT before imaging using a C-digit blot scanner (LI-COR, Lincoln, NE, USA).

### RNA quantification

Total RNA was extracted with QIAzol Lysis Reagent and then purified using a miRNeasy Mini kit according to the manufacturer’s protocol (Qiagen, Germantown, MD, USA). After purification, the RNA concentration was quantified using a Qubit microRNA Assay Kit at excitation/emission wavelengths of 500/525 nm.

**Supplementary Figure 1.**
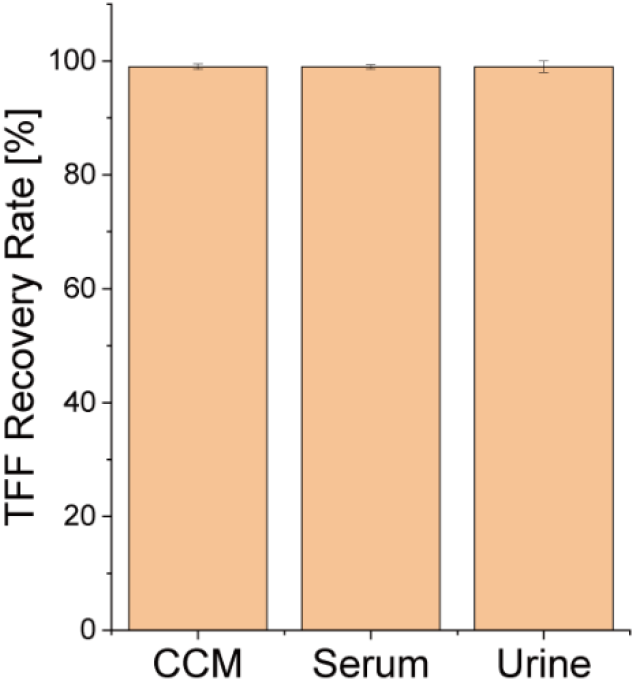
EV recovery rate after TFF processing. Cell culture medium (n = 5), serum (n = 5), and urine (n = 5) were purified using TFF and enriched into a volume of 2 mL.

**Supplementary Figure 2.**
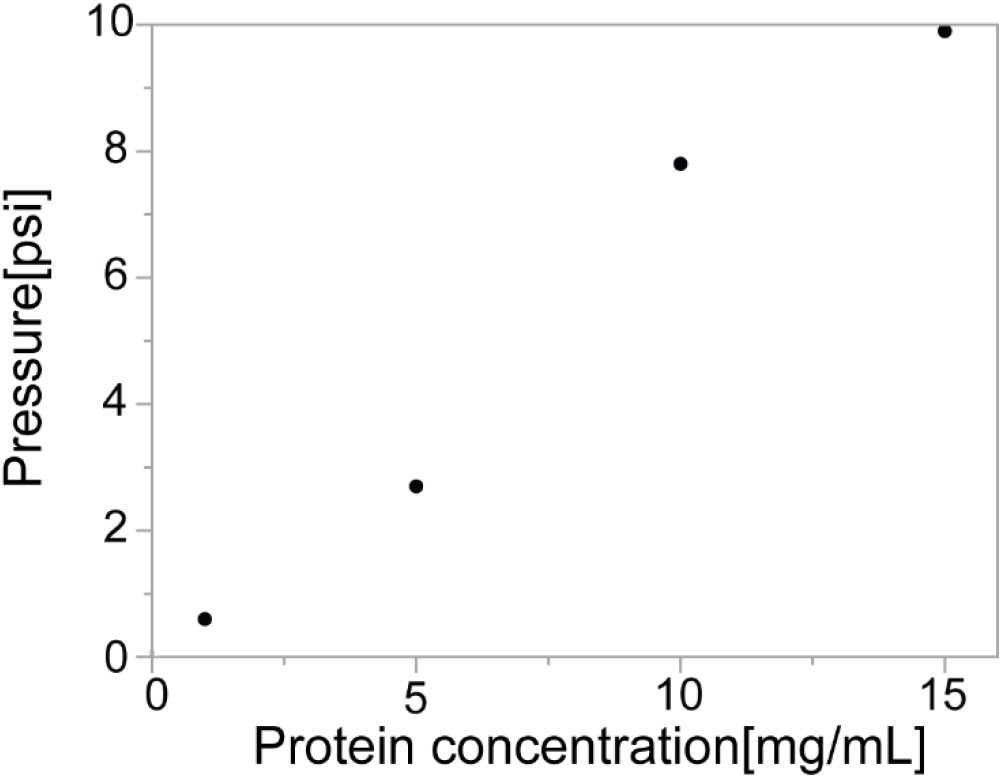
Pressure in the TFF stage of iSUF when processing bovine serum albumin (BSA) solutions at different concentrations using a fixed flow rate of 35 mL/min. 15 mg/mL was the maximum protein concentration in the TFF system to maintain the system pressure below 10 psig.

**Supplementary Figure 3.**
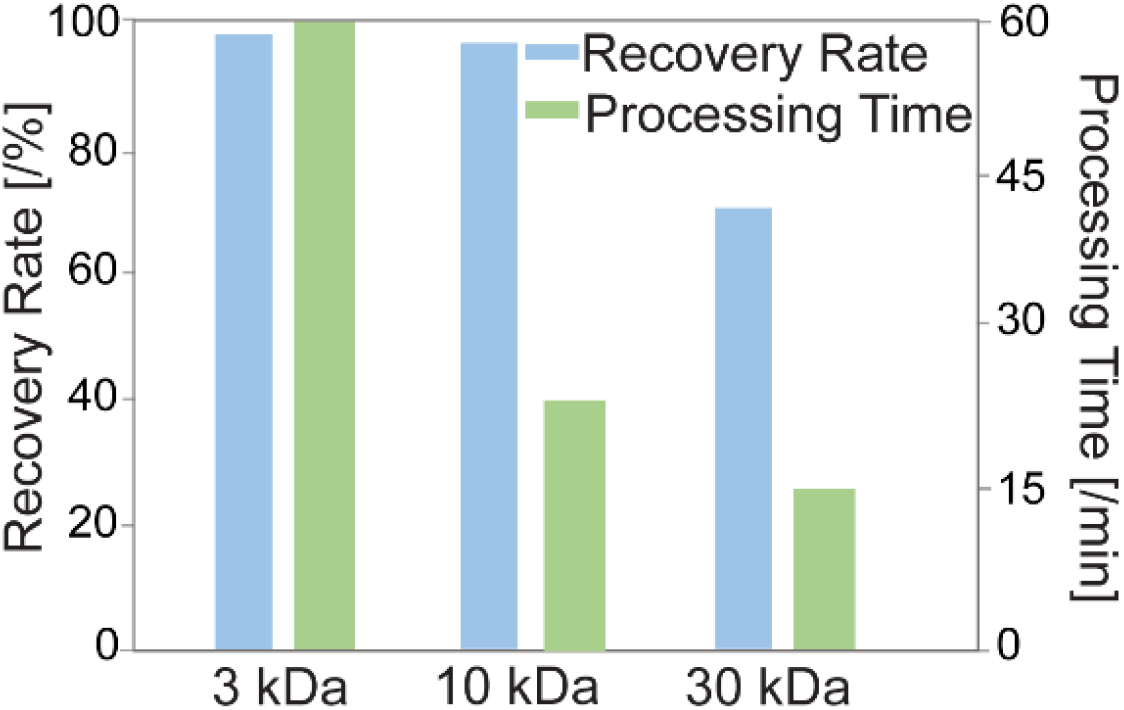
EV recovery rate and processing time of centrifugal units (iSUF, stage 2) with different MWCOs. 3 kDa, 10 kDa, and 30 kDa MWCO centrifugal units obtained over 99%, 95%, and 70% recovery rate, respectively. They took 60 min (3 kDa MWCO), 20 min (10 kDa MWCO), and 15 min (30 kDa MWCO) to spin down the sample to a final volume of 100 μL.

**Supplementary Figure 4.**
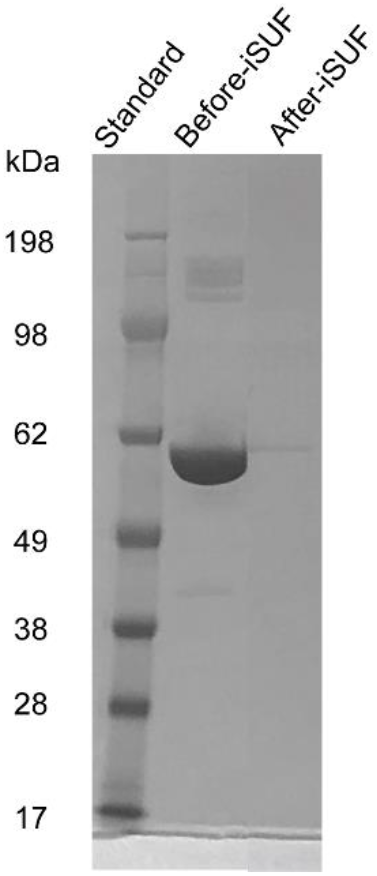
SDS-PAGE of 10% BSA solution before and after iSUF processing. BSA was extensively removed after iSUF processing.

**Supplementary Figure 5.**
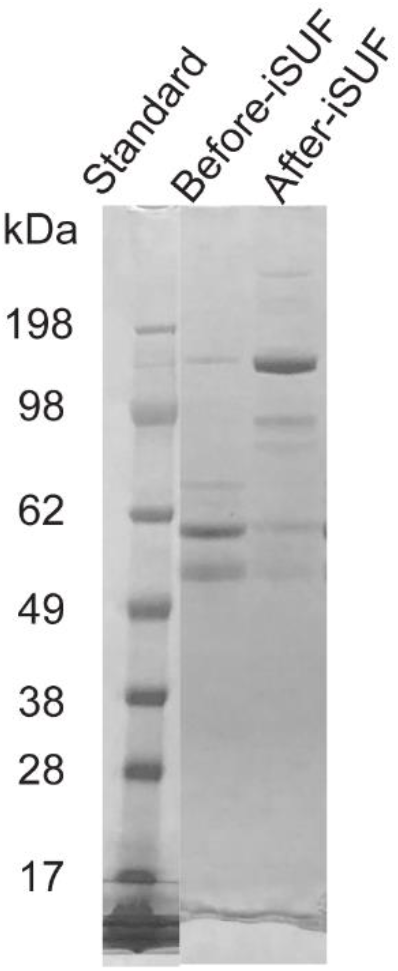
SDS-PAGE of CCM before and after iSUF processing. BSA was extensively removed after iSUF processing.

**Supplementary Figure 6.**
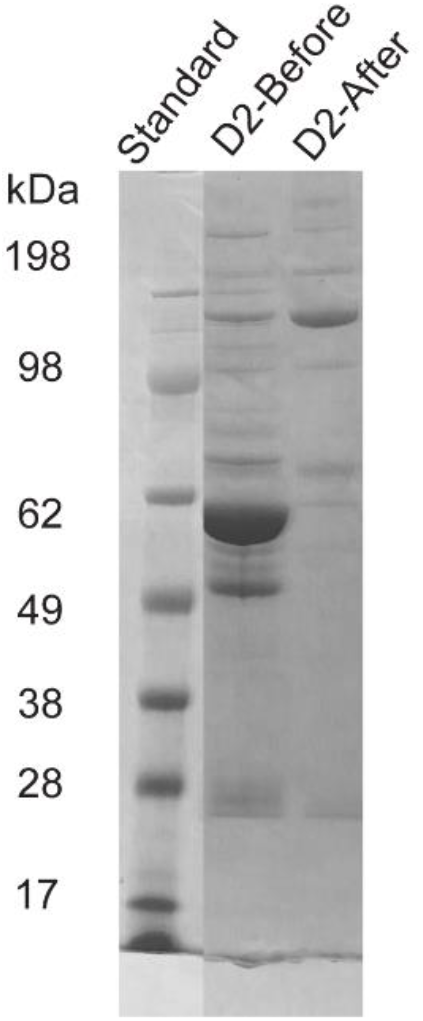
SDS-PAGE of serum samples before and after iSUF processing. HSA was extensively removed after iSUF processing.

**Supplementary Figure 7.**
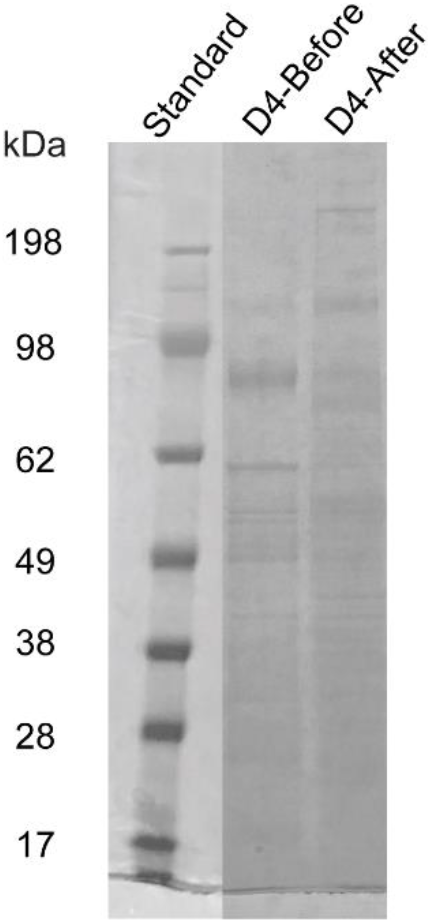
SDS-PAGE of urine samples before and after iSUF processing. HSA was extensively removed after iSUF processing.

**Supplementary Figure 8.**
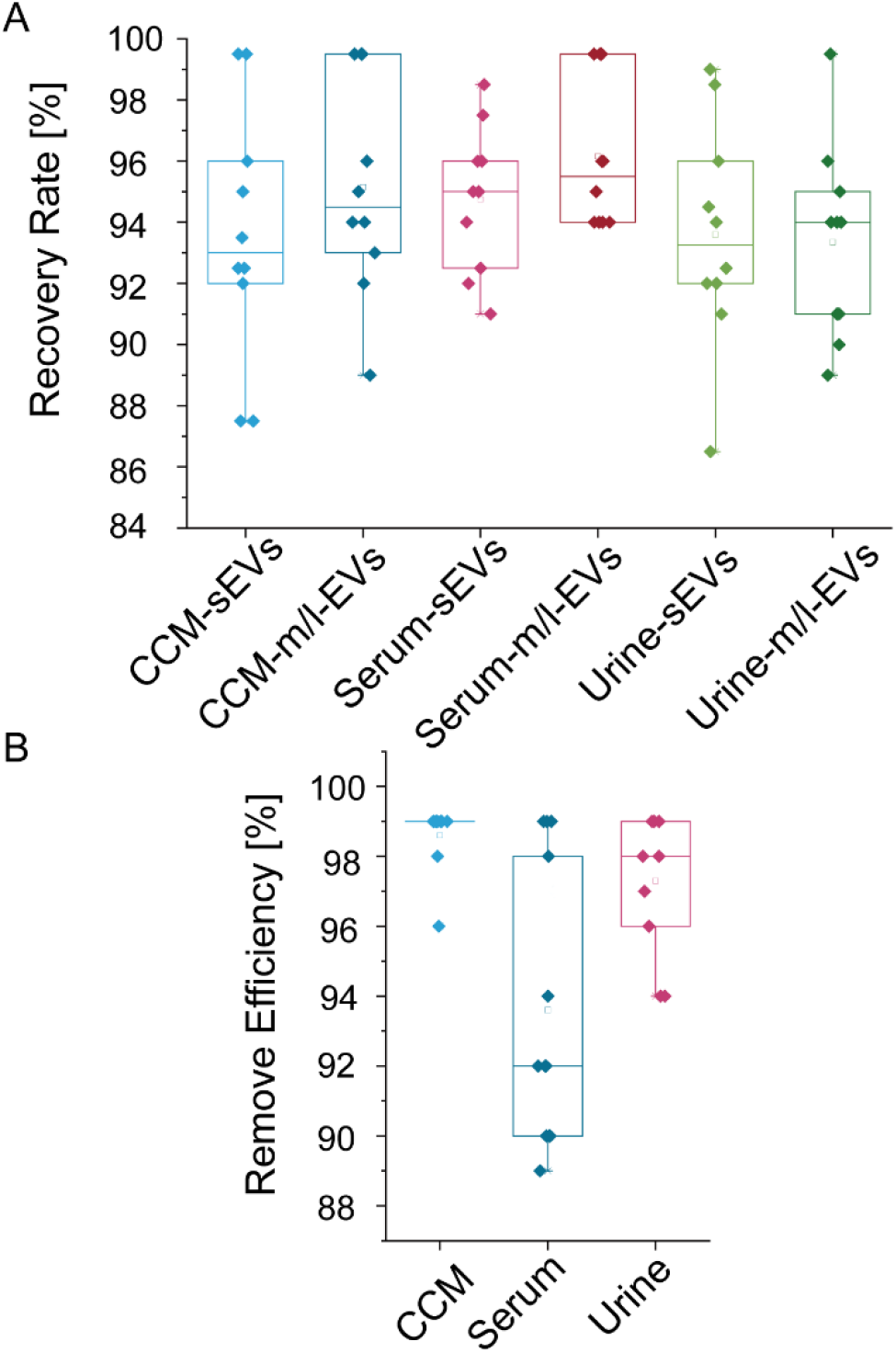
**A)**. Repeatability evaluation of EV recovery rate for samples processed by iSUF. Cell culture medium (n = 10), serum (n = 10), and urine (n =10) were purified by iSUF and enriched into a final volume of 100 μL. **B)**. Repeatability evaluation of protein removal efficiency for samples processed by iSUF.

**Supplementary Figure 9.**
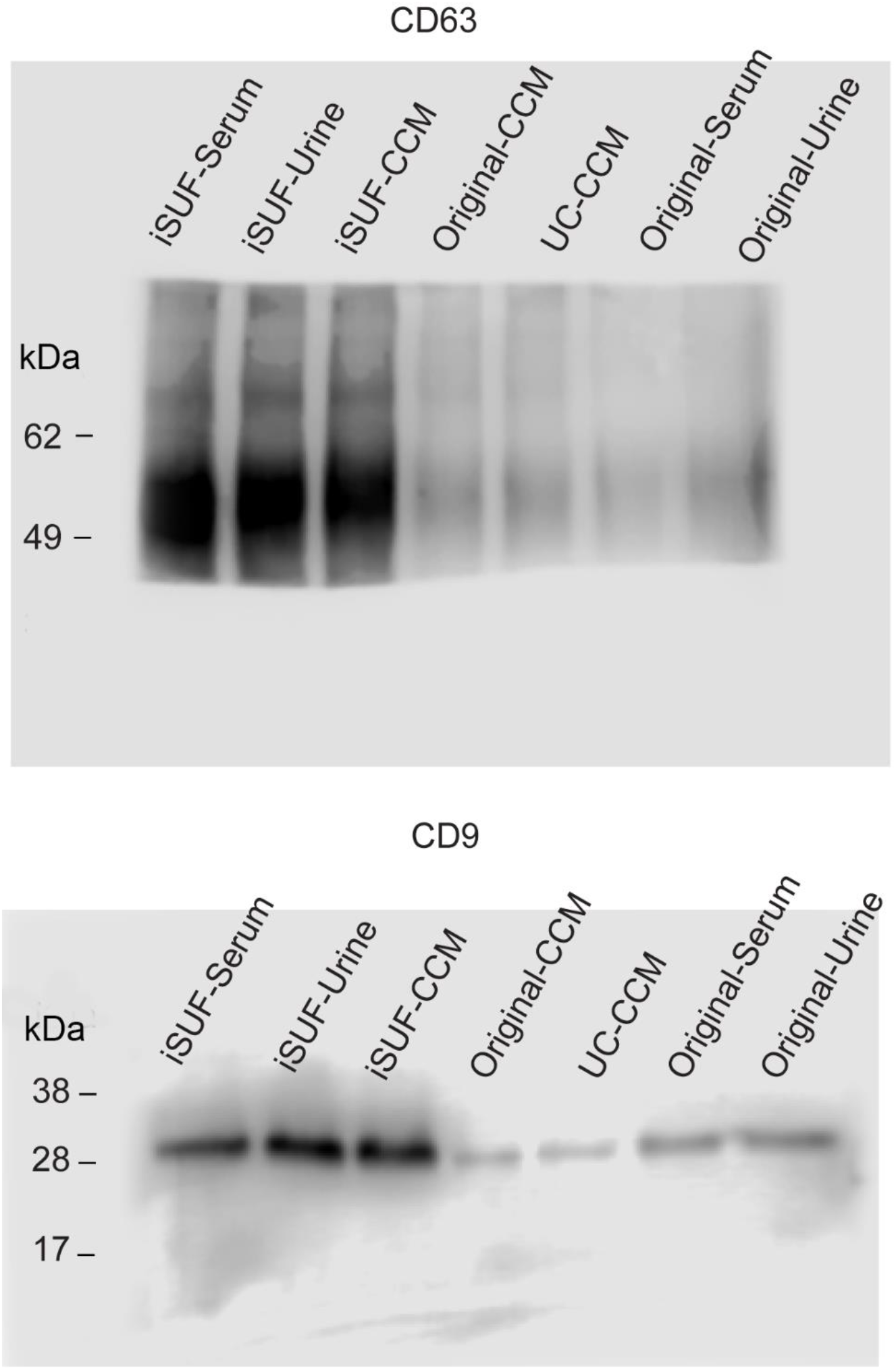
Full-length western blots for cropped images in Fig. 4E.

**Supplementary Figure 10.**
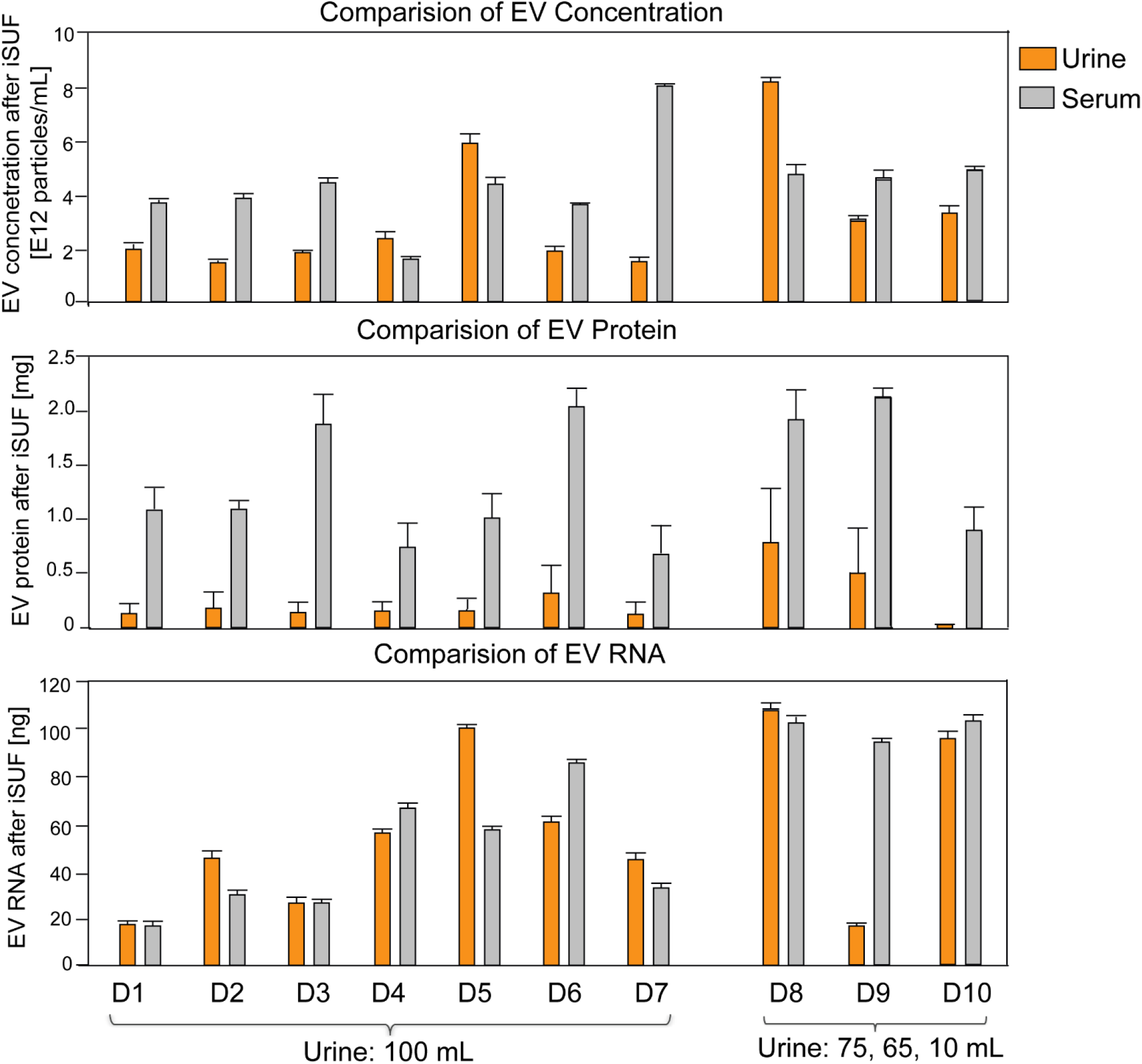
Comparison of EV concentration, protein, and RNA content in iSUF-urine and iSUF-serum samples from seven healthy donors. The EV concentration and RNA content showed comparable values between urine and serum (n = 10; p > 0.05), while protein content in urine was 10 times lower than in serum.

**Supplementary Figure 11.**
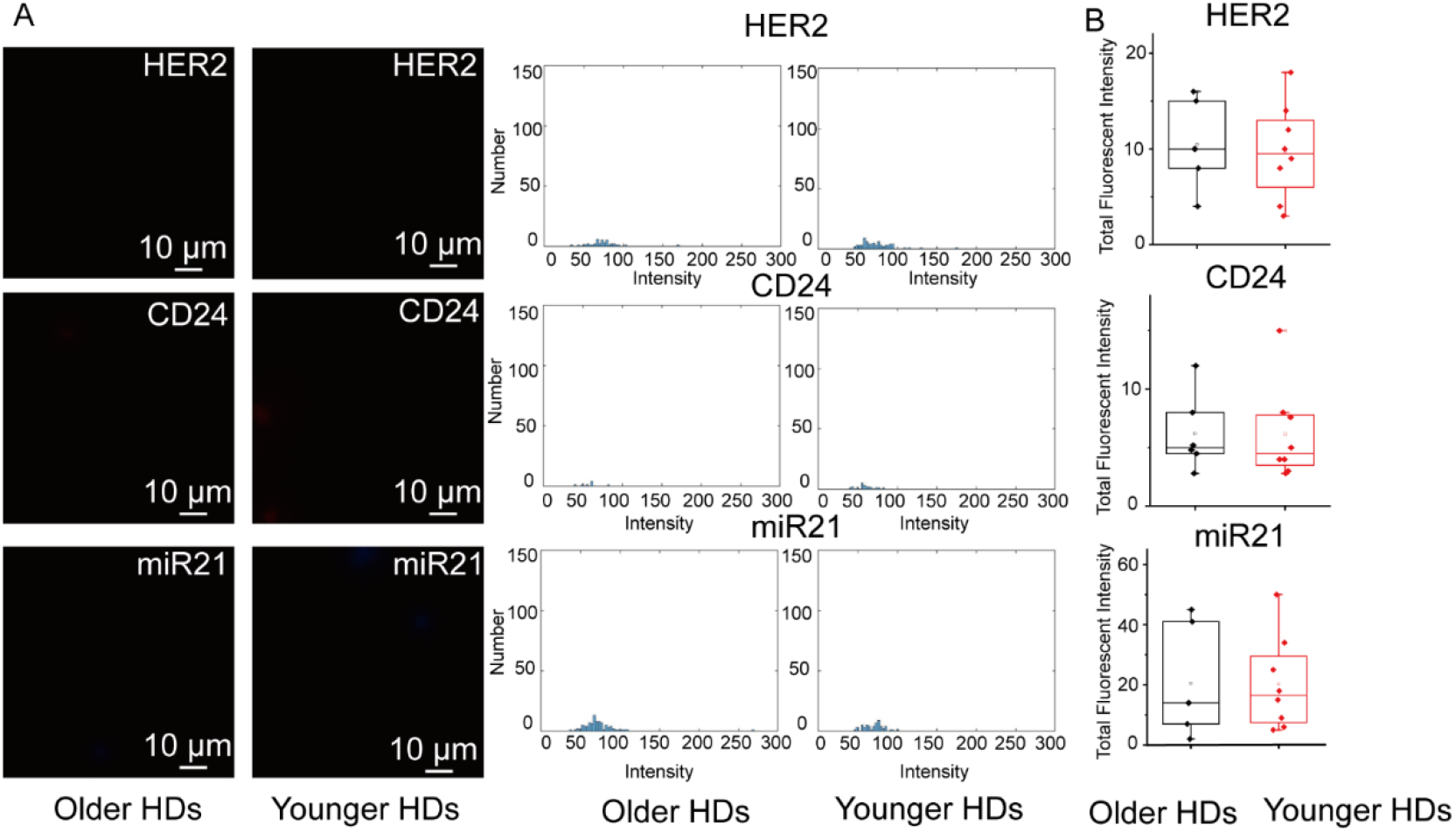
Levels of expression of BC biomarkers in young and old healthy donor serum samples. For older HDs, the mean age was 58.7 ± 5.6 years (SD); for younger HDs, the mean age was 24.8 ± 2.1 years (SD). There was not a statistically significant difference in fluorescence signal for both cohorts (p > 0.05). **A)** Characteristic fluorescence images of HER2, CD24, miR21. **B)** Total fluorescence intensity quantification of HER2, CD24, miR21.

**Supplementary Figure 12.**
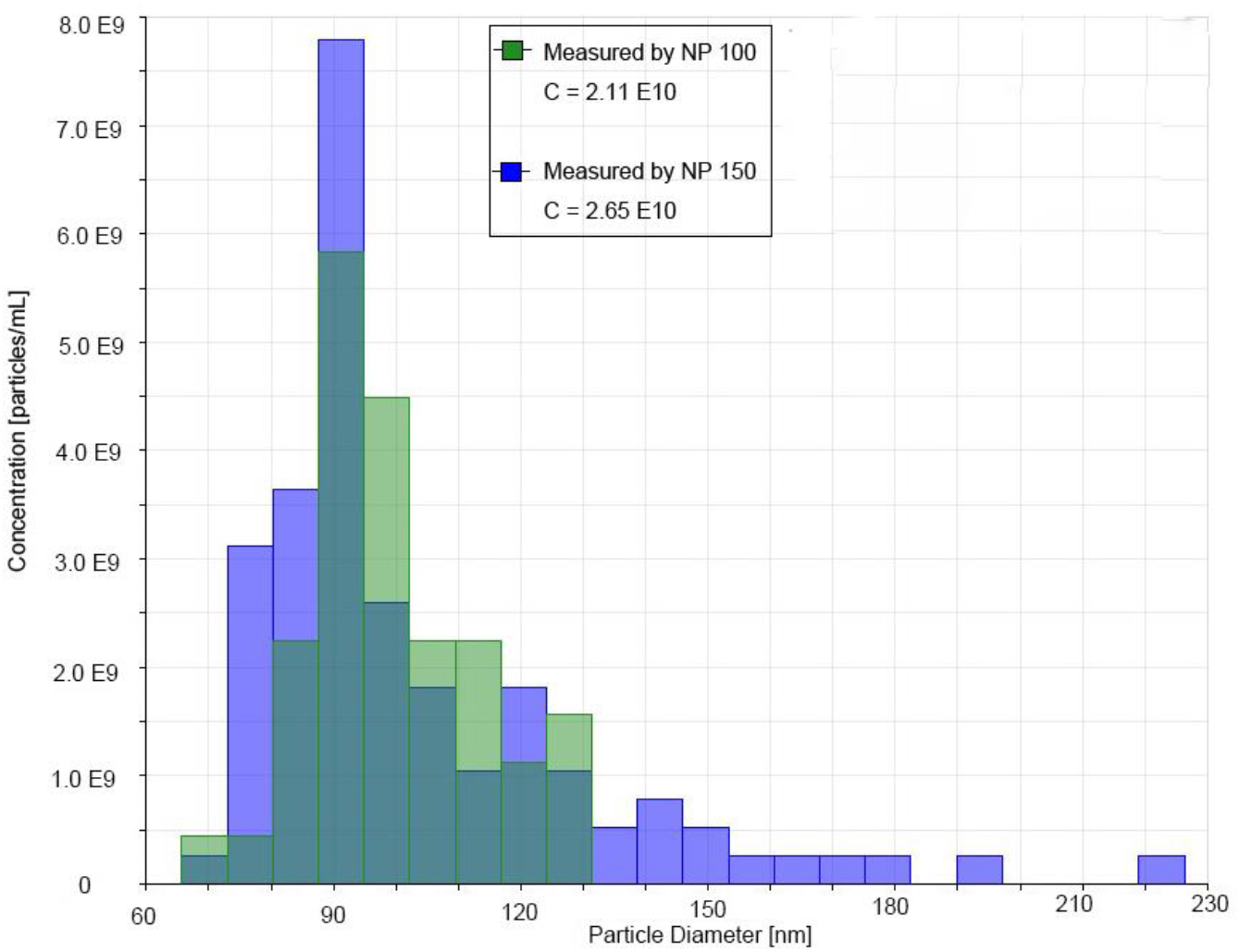
Size distribution of EVs in a CCM sample processed by iSUF and measured by a TRPS method (qNano) using an NP100 and NP150 stretchable membrane.

**Supplementary Figure 13.**
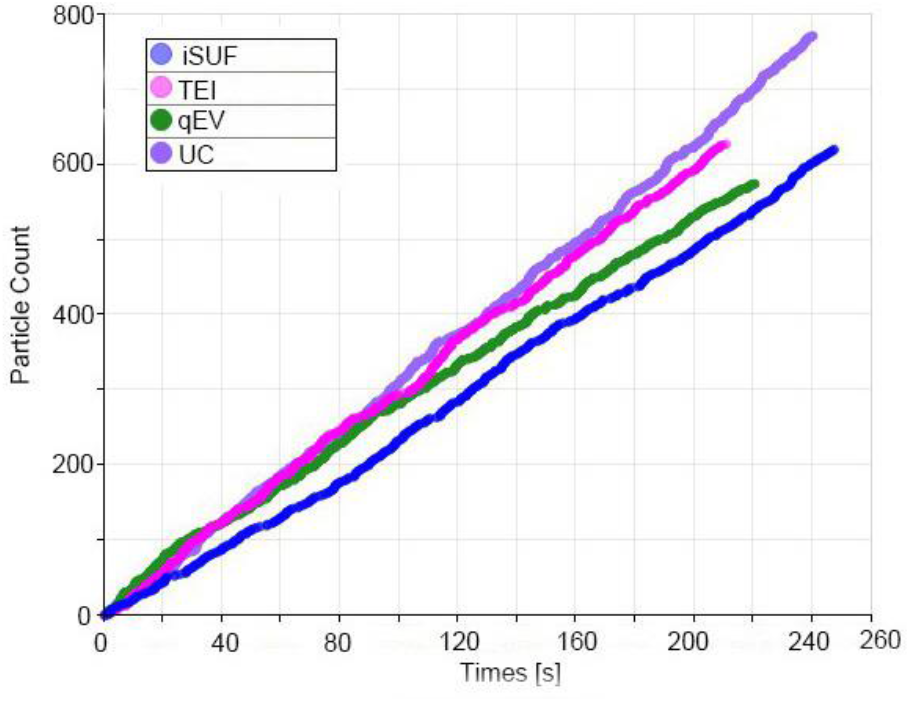
Particle count plot of EVs isolated by iSUF, TEI, qEV, and UC. The graph indicates an overall constant particle detection.

**Supplementary Table 1.** Comparison of 300 and 500 kDa TFF filter performance on CCM, urine, and serum.

| TFF filter<br>MWCO | CCM (50 mL) |  | Urine (100 mL) |  | Serum (0.5 mL) |  |
| --- | --- | --- | --- | --- | --- | --- |
|  | Protein<br>removal<br>efficiency | Processing<br>time | Protein<br>removal<br>efficiency | Processing<br>time | Protein<br>removal<br>efficiency | Processing<br>time |
| 300 kDa | >78% | 150 min | >80% | 200 min | >75% | 300 min |
| 500 kDa | up to 99% | 80 min | up to 99% | 100 min | up to 99% | 120 min |

**Supplementary Table 2.** Protein concentration of the TFF enrichment step.

|  | U251 Supernatant | Urine | Serum |
| --- | --- | --- | --- |
| Initial protein concentration (mg/mL) | $1.20 \pm 0.13$ (in 50 mL) | $1.40 \pm 0.50$ (in 10~100 mL) | $10.0 \pm 2.0$ (dilute 0.5 mL of serum to total volume of 7mL in PBS) |
| Protein concentration after TFF enrichment step (mg/mL) | $2.30 \pm 0.38$ | $1.90 \pm 0.31$ | Not applied |

**Supplementary Table 3.** EV concentration, purity, and RNA content comparison among different EV purification platforms.

| | Concentration<br>(particles/mL) | Free protein<br>(μg/mL) | Purity<br>(particles/μg) | RNA (ng) |
| --- | --- | --- | --- | --- |
| qEV | 8.19E10±2.62E10 | 15.39±4.06 | 5.94E09±3.08E09 | 8.29±7.58 |
| UC | 7.46E10±2.25E10 | 11841.0±2375.6 | 6.77E06±3.30E06 | 5.52±5.02 |
| TEI | 5.70E11±2.79E11 | 67.2±29.8 | 1.32E10±1.43E10 | 15.23±8.24 |
| iSUF | 4.15E12±4.66E11 | 6.32±4.28 | 1.32E12±1.26E12 | 53.72±41.02 |

**Supplementary Table 4.**
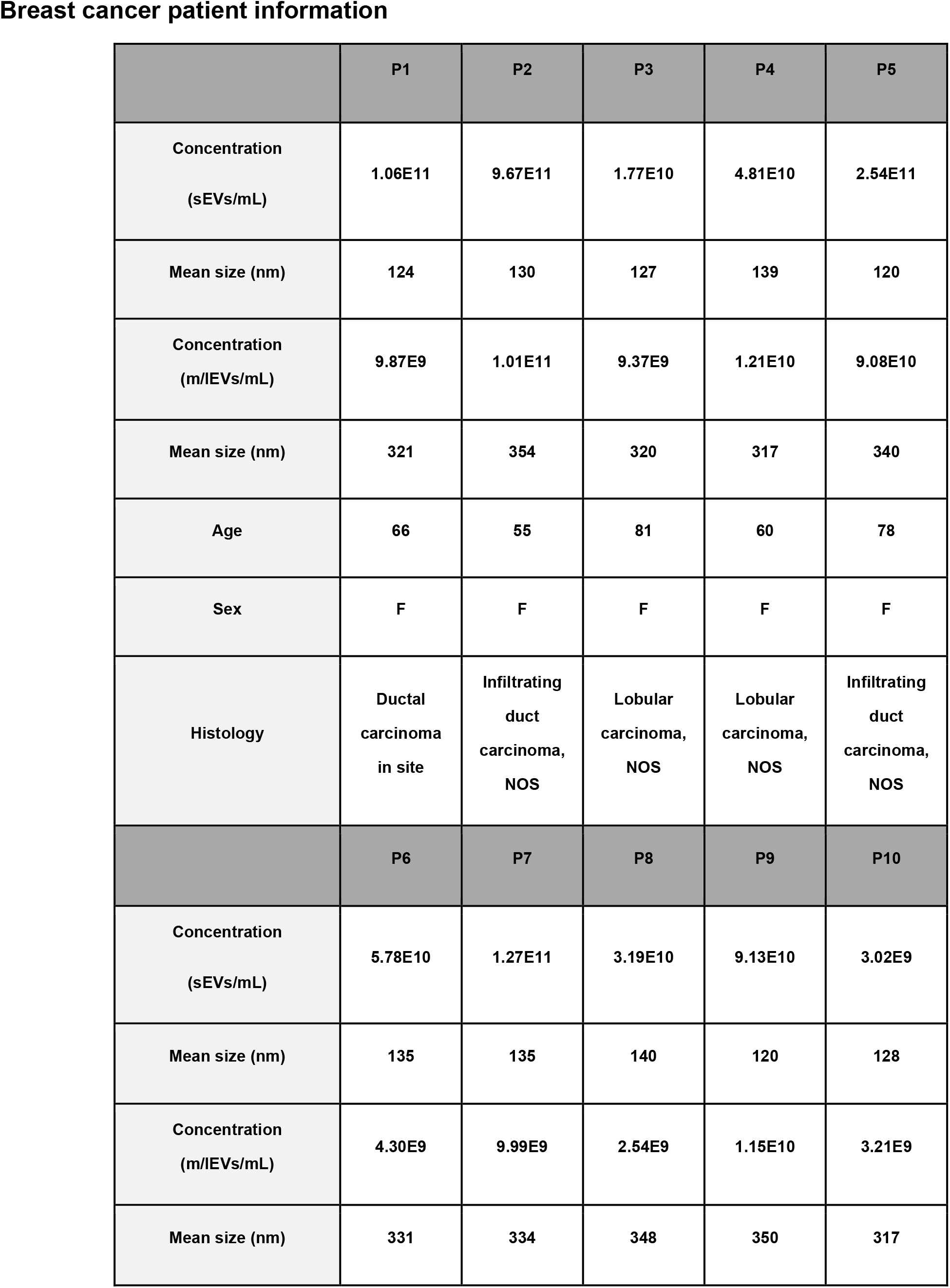
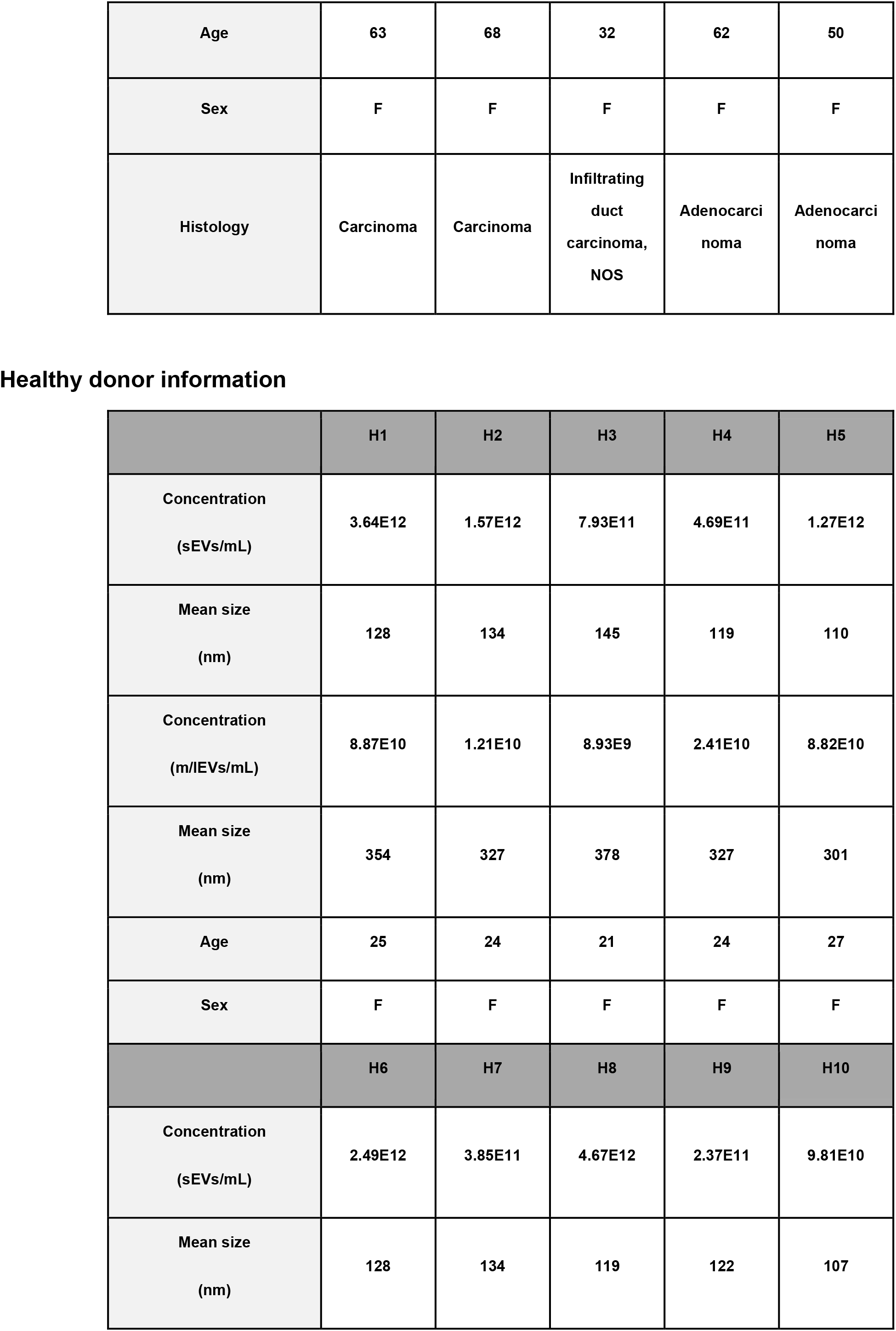
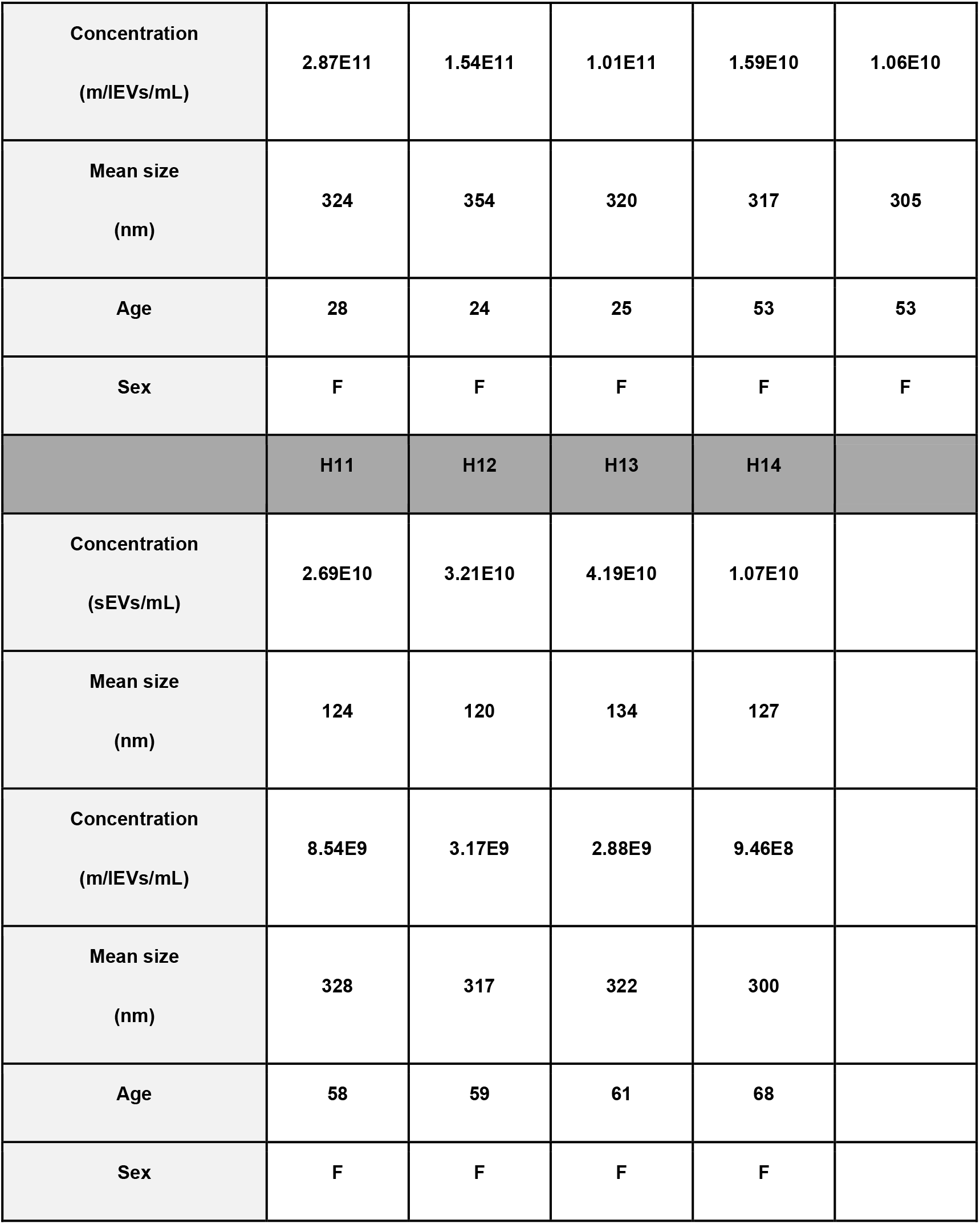

